# A genotoxin associated with colorectal cancer linked to gut dysbiosis in children with cystic fibrosis

**DOI:** 10.1101/2025.09.04.674286

**Authors:** Kaitlyn E. Barrack, Sarvesh V. Surve, Ana V. de Sousa Bezerra, Caitlin E. Murphy, Shannon M. Soucy, Miguel A. Aguilar Ramos, Rebecca A. Valls, Rebekah D. Ruff, Emily P. Balskus, Julie L. Sanville, Juliette C. Madan, George A. O’Toole

**Affiliations:** Department of Microbiology and Immunology, Geisel School of Medicine at Dartmouth, Hanover, New Hampshire, USA; Department of Biomedical Data Science, Geisel School of Medicine at Dartmouth, Hanover, New Hampshire, USA; Department of Chemistry and Chemical Biology, Harvard University, Cambridge, MA, USA; Howard Hughes Medical Institute, Harvard University, Cambridge, MA, USA; Division of Pediatric Gastroenterology, Department of Pediatrics, Dartmouth Hitchcock Medical Center, Lebanon, New Hampshire, USA; Departments of Psychiatry and Pediatrics, Dartmouth Hitchcock Medical Center, Lebanon, New Hampshire, USA; Geisel School of Medicine at Dartmouth, Hanover, New Hampshire, USA

**Keywords:** cystic fibrosis, gut microbiome, colibactin, dysbiosis, physiology

## Abstract

Cystic fibrosis (CF) substantially alters the gastrointestinal microbiome from an early age, leading to significant changes in microbial composition and functionality. This study explores the physiological and microbiological factors contributing to dysbiosis in children with cystic fibrosis (cwCF), characterized by an increase in potentially pathogenic *Escherichia coli* and a decrease in beneficial anaerobes such as *Bacteroides*. In this study, we employed an in vitro medium representative of the nutritional environment of the CF colon to test the role of factors including mucin, fat, bile, pH, antibiotics and features associated with inflammation (e.g., nitrate, sulfate, formate, reactive oxygen species) on growth of clinical isolates of *E. coli* and *Bacteroides* spp. We further examined interactions between these two microbes under CF-like conditions to understand modulators of microbial competition, and identified glycerol, a surrogate of increased fat, as a significant driver of altered microbial competition. Finally, we investigated genetic determinants influencing these microbial interactions, with the focus on glycerol metabolism, by performing a transposon mutagenesis screen in *E. coli*. Results of this screen pointed to the role of colibactin production in mediating this microbial competition; colibactin is a DNA-damaging genotoxin associated with the increased risk of colorectal cancer (CRC) in CF populations. This work enhances our understanding of mechanisms of microbial competition in the CF gut, while potentially enhancing our understanding of colorectal cancer risk in persons with CF through the identification of early-life microbial biomarkers.

**Significance Statement:** The risk of CRC development in CF populations is significantly increased. This study examines the interplay of altered intestinal physiology in the microbial dysbiosis common in the CF gut, implicating the high fat environment in a competition-mediated depletion of immune-modulating *Bacteroides*. This work identifies candidate features of the young CF intestine and gut microbiome that may contribute to advanced development of CRC in these populations, informing potential therapeutic approaches.

## Introduction

The cystic fibrosis (CF) gastrointestinal tract is altered both physiologically and microbiologically from a young age [1-6]. Key taxonomical shifts observed in CF cohorts include increased Proteobacteria, largely driven by *Escherichia coli* [3-7], and decreased anaerobes that produce short chain fatty acids (SCFAs), such as *Bacteroides, Akkermansia, Faecalibacterium* and *Roseburia* [3, 6, 8]. These shifts are reported as early as 6 weeks of age in children with CF (cwCF) [1]. Many factors likely contribute to this microbial imbalance, with large contributions from altered physiology [9], including nutrition [10-12], antibiotic use [13] and microbial competition. However, the role of such factors has not been thoroughly investigated in the context of CF.

Previous research investigated the role of increased fat in the CF intestine, driven by diet and fat malabsorption [14, 15], on enrichment of *E. coli* [7]. *E. coli* isolates originating from a CF donor were better adapted to growth conditions supplemented with glycerol, a surrogate for increased fat, than isolates originating from a nonCF donor [7]. CF isolates not only grow to a greater extent on glycerol, but the transcriptome of these isolates exhibited a decreased stress response to glycerol as a sole carbon source compared to nonCF isolates. Rather than these isolates upregulating glycerol-utilization genes, which would be expected, the CF isolates seem to lose the signal for growth inhibition and stress response that glycerol typically induces [7]. These findings implicate the role of fat and its breakdown products in shifting microbial abundance and function, while driving adaptation to altered physiology in the context of CF.

A dysbiotic intestinal environment and microbial composition modulates functionality in CF [16, 17], delays maturation [3, 6], and contributes to increased inflammation [18]. From early ages, cwCF have increased markers of inflammation [17, 19], which correlate with microbial markers of dysbiosis, namely increased *E. coli* [19]. Importantly, gut dysbiosis and inflammation are implicated in the development of colorectal cancer (CRC); persons with CF (pwCF) have 10 times the risk of developing CRC compared to the general population [20, 21]. Mechanisms underlying this increased risk in CF remain undetermined. In nonCF populations, pathogenic *E. coli* strains capable of producing the genotoxin colibactin, which promotes DNA damage, via expression of a polyketide synthesis (*pks*) island have been linked with increased CRC risk, microbial dysbiosis and inflammation [22-25]. However, limited research has investigated the role of colibactin in CF populations, and there is inconclusive evidence of increased prevalence of *E. coli pks+* strains for pwCF [26, 27]. Thus, determining physiological and/or microbiological biomarkers, especially in early life, is imperative to understanding mechanisms of CRC development and improving therapeutic potential in CF.

In this study, we first aimed to identify physiological features of the CF intestine that may be involved in driving microbial dysbiosis, namely increased *E. coli* and decreased *Bacteroides*, using a previously developed in vitro medium that models the CF gut [28]. We also investigated the dynamics of the microbial interaction of these species under various CF-like physiologies, highlighting an important role for glycerol in this microbial interaction. Finally, we probed the genetic components of *E. coli* involved in modulating the microbial interaction with *Bacteroides*, identified genes whose products are involved in glycerol metabolism and colibactin synthesis, and performed a characterization of their prevalence and function in the colibactin-encoding *pks* island CF and nonCF *E. coli* isolates and stool metagenomes. Results suggest an increase in *pks* gene prevalence and abundance in CF stool metagenomes, compared to nonCF stool metagenomes, coupled with a trend for increased colibactin production in CF stool. Our work identifies microbial and physiological factors in young children with CF potentially contributing to the development of CRC later in life.

## Results

### *Bacteroides vulgatus* is sensitive to CF-relevant physiological features and to *E. coli* competition

To test the hypothesis that physiological features relevant to the CF intestine may be driving taxonomical shifts, we decided to focus on two key microbes that are consistently and significantly differentially abundant in CF cohorts: *Escherichia coli* and *Bacteroides* species (spp.). Metagenomic studies have specifically identified *B. vulgatus* (recently renamed *Phocaeicola vulgatus* but referred to as *B. vulgatus* in this study) as being depleted in CF stool samples [19]. We grew CF and nonCF clinical isolates of *B. fragilis, B. thetaiotaomicron, B. vulgatus* and *E. coli* (**Supplemental Table 1**) in a recently developed CF intestinal medium called CF-MiPro [28]. This medium was designed and validated to mimic the nutritional environment of the CF colon, including excess fats, mucin, bile, reactive oxygen species (ROS), alternative nutrients associated with inflammation (e.g., nitrate, sulfate, formate), lower pH, and sub-lethal levels of antibiotics. CF-MiPro was developed using MiPro, a medium that mimics the healthy gut environment as the foundational medium base [29], then the aforementioned features were supplemented at a gradient of concentrations to represent low and median levels detected in reported CF stool samples [28].

We hypothesized that clinical isolates of *E. coli* would have a growth enhancement in CF-MiPro, while *B. vulgatus* clinical isolates grown in monoculture would be susceptible to this in vitro environment, mirroring what is seen in CF gut microbiome samples [19]. Indeed, after 24h growth in MiPro, low-CF-MiPro and median-CF-MiPro, there is an observed dose-dependent depletion of all tested *B. vulgatus* clinical isolates (**Figure 1A**). This depletion is not observed in the tested *B. fragilis* nor *B. thetaiotaomicron* strains, suggesting increased tolerance to CF-MiPro and thus a CF-like intestinal environment for these species. Contrary to our hypothesis, *E. coli* clinical isolates do not exhibit a growth enhancement in CF-MiPro, but rather similar colony forming units per milliliter (CFU/mL) across all conditions. We observed a trend towards reduced viability in median-CF-MiPro, but this difference was not significant.

**Figure 1.**
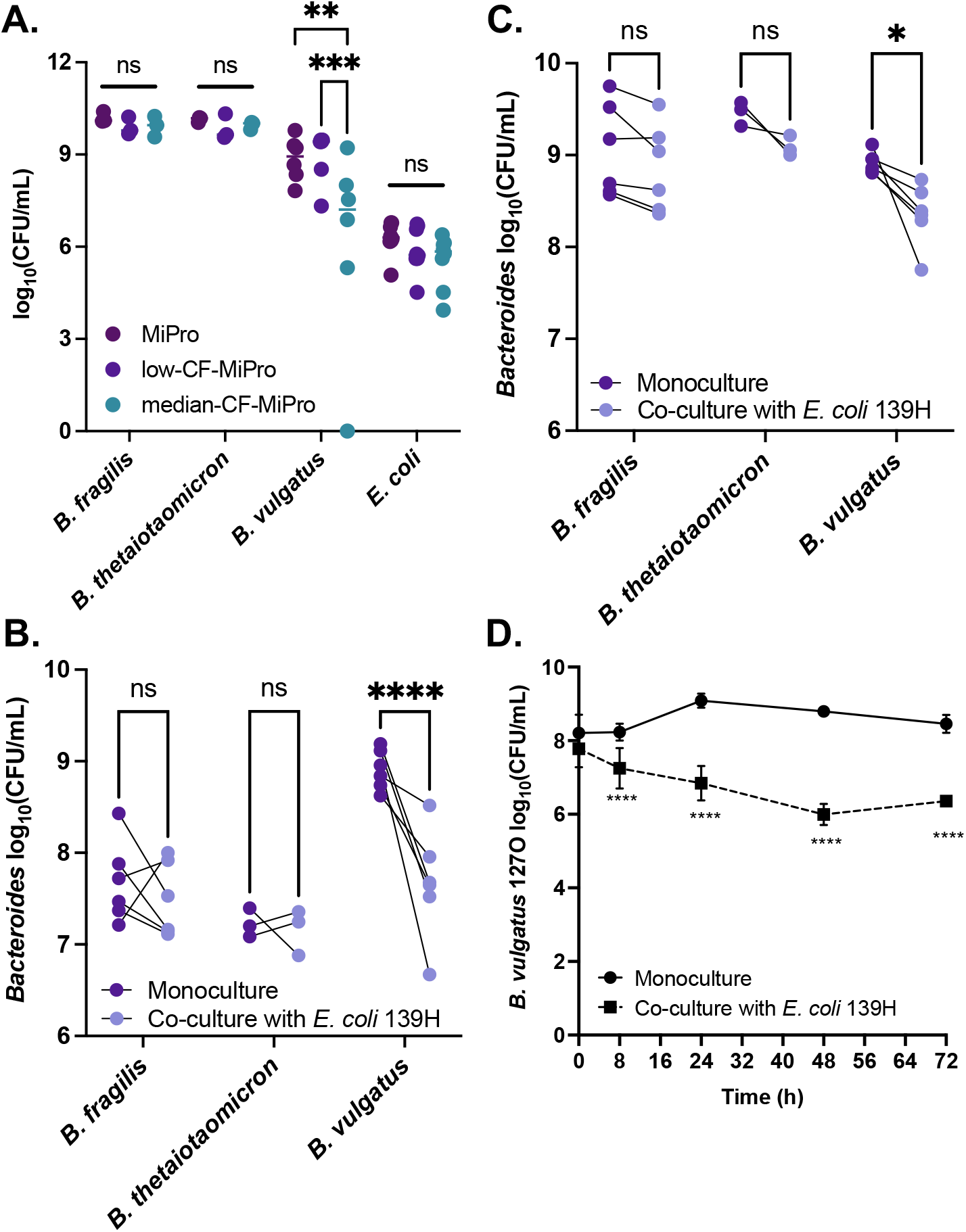
*Bacteroides vulgatus* is susceptible to CF-like growth conditions and competition with *E. coli*. **(A)** Clinical isolates of *Bacteroides fragilis* (n=3), *B. thetaiotaomicron* (n=3), *B. vulgatus* (n=5), and *E. coli* (n=6) were grown in monoculture in MiPro (dark purple), low-CF-MiPro (purple), and median-CF-MiPro (teal) for 24h, anaerobically. Growth is quantified via log_10_(CFU/mL). **(B-C)** Clinical isolates of *B. fragilis* (n=6), *B. thetaiotaomicron* (n=3) and *B. vulgatus* (n=5) were co-cultured anaerobically with a CF clinical isolate of *E. coli*, 139H, in a 1:1 ratio in **(B)** BHIS pH 7 or **(C)** MiPro pH 7 for 72h. Growth of each respective *Bacteroides* strain is displayed as log_10_(CFU/mL) on the y-axis when in monoculture (dark purple) or co-culture with *E. coli* (light purple), with a line connecting each strain. **(D)** Growth kinetics of one *B. vulgatus* strain, 127O, in co-culture with *E. coli* 139H in MiPro pH 7 for 72h. CFU/mL were plated at 0h, 8h, 24h, 48h and 72h. Statistical analysis performed using 2-way ANOVA with Šídák’s multiple comparisons (^*^, p < 0.05; ^**^, p < 0.01; ^***^, p < 0.005; ^****^, p < 0.001).

Next, we investigated the role of microbial competition between these two genera, since altered microbial interactions likely occur in the CF intestinal lumen [30] and may exacerbate the observed dysbiosis. Here, we inoculated a CF isolate of *E. coli*, 139H, in a 1:1 ratio with strains of *B. fragilis, B. thetaiotaomicron* and *B. vulgatus* and co-cultured each pair of isolates for 72h in anoxic conditions. To address the impact of competition with *E. coli* on *Bacteroides* growth, we used *Bacteroides* grown in monoculture (assessed by CFU/mL) as a positive control. In two separate rich media conditions buffered to pH 7, supplemented brain-heart infusion broth (BHIS) and MiPro, the viability of *B. vulgatus* strains was significantly reduced in the presence of *E. coli* 139H (**Figures 1B and C**). This reduction was not observed in *B. fragilis* nor *B. thetaiotaomicron* co-cultures, although there was a trend towards reduced viability for both species. We confirmed that the reduction of *B. vulgatus* viability occurs independently of enhanced *E. coli* growth - *E. coli* CFU/mL in co-culture is not significantly increased compared to its own monoculture controls (**Supplemental Figure 1A**).

To observe the timing at which this reduction of *B. vulgatus* viability by *E. coli* occurs, we performed a growth kinetics assay with CF isolate *B. vulgatus* 127O and *E. coli* 139H in MiPro pH 7 for 72h, collecting CFU/mL of each strain at 8h, 24h, 48h, 72h. Reduction of the viability of 127O is observed as early as 8h in co-culture with *E. coli*, compared to monoculture controls (**Figure 1D**). Again, this reduction in viability is largely independent of enhanced *E. coli* growth, whose viable counts are stable or slightly decreased at early time points, in co-culture with *B. vulgatus* compared to monoculture controls (**Supplemental Figure 1B**). These results suggest that baseline competition occurs between these two microbes under in vitro conditions that mimic the healthy intestine. Next, we aimed to explore how this interaction was mediated in CF-like growth conditions.

### Linear regression identifies bile and glycerol as key contributors to the reduction of *B. vulgatus* viability in the context of the CF intestinal environment

To address the question of how CF-relevant physiological features impact growth and competition of intestinal flora, we prepared MiPro medium with individual components altered to the concentration used in CF-MiPro, then grew clinical isolates of *B. vulgatus* (n=6) and *E. coli* (n=8) in mono- and co-culture for 72h. The components that were tested include: MiPro pH 7 (control), MiPro pH 6, or MiPro pH 7 individually supplemented with mucin, bile salts, glycerol, sodium nitrate, sodium sulfate, sodium formate, hydrogen peroxide or Bactrim, an antibiotic commonly used in CF clinics. See the **Materials and Methods** for respective concentrations of each component. After 72h of growth in each respective CF-relevant feature, CFU/mL were calculated for each species and associated metadata were assigned (e.g., species, strain, monoculture vs. co-culture, strain identification of co-culture partner, feature and respective concentration, log-transformed CFU/mL; **Supplemental Table 2**). A linear model was performed to analyze the contribution of each feature on CFU/mL of each species.

Unsurprisingly, species and microbial competition (monoculture vs. co-culture) each play a significant role on overall CFU/mL (**Supplemental Table 3**). Species identification contributes to 3.26% of variance in the model (P<0.001) and competition status accounts for 0.148% of the model (P=0.007). To account for these contributions and accurately address which physiological features impact each species under each competition status, we subset the metadata by species and co-culture, then performed separate linear models for each category: *E. coli* in monoculture (**Supplemental Table 4**), *E. coli* in co-culture (**Supplemental Table 5**), *B. vulgatus* in monoculture (**Supplemental Table 6**), and *B. vulgatus* in co-culture (**Supplemental Table 7**).

*E. coli* survival in monoculture is significantly increased by numerous CF-like physiological features, including mucin (P=0.003), bile (P=0.001), glycerol (P=0.002), nitrate (P=0.003), sulfate (P=0.007) and formate (P=0.014) compared to the MiPro pH 7 control (**Supplemental Table 4**). All tested features account for 17.8% of variance in the model. Notably, the degree of response, or change in CFU/mL, of each feature (as represented through the “Estimate” parameter) is minimal (less than 1), suggesting that while that changes we note are significant, the shifts in survival are modest across *E. coli* strains. There is some overlap in features that significantly affect *E. coli* survival in co-culture with *B. vulgatus*, as well. Features that direct survival in both mono- and co-culture include bile (P=0.049) and glycerol (P=0.001) (**Supplemental Table 5**). Additional features that impact co-culture growth of *E. coli* include hydrogen peroxide (P<0.001), pH 6 (P<0.001) and antibiotics at low-CF-MiPro (P<0.001) and median-CF-MiPro concentrations (P<0.001). Similar to what is observed in monoculture, bile and glycerol increase *E. coli* CFU/mL in co-culture, although the Estimate parameters indicate very minimal shifts (0.03 and 0.06, respectively). However, hydrogen peroxide (-0.26), lower pH (-0.3) and antibiotics (low: -0.33, median: -0.31) all decrease *E. coli* CFU/mL in co-culture, relative to MiPro pH 7 control. All tested features account for 26.6% of variance in the model.

*B. vulgatus* survival in CF-like conditions is significantly impacted by fewer features, compared to *E. coli*. In monoculture, only bile at median concentrations has a significant negative impact on survival (Estimate = -5.17, P<0.001, **Figure 2A, Supplemental Table 6**). All tested features account for 26.2% of variation in the model. In co-culture with *E. coli*, bile (Estimate = -3.36, P<0.001) and glycerol (Estimate = -3.15, P<0.001) are negatively associated with *B. vulgatus* survival (**Figure 2A, Supplemental Table 7**). All tested features account for 14.4% of variation in the co-culture model.

**Figure 2.**
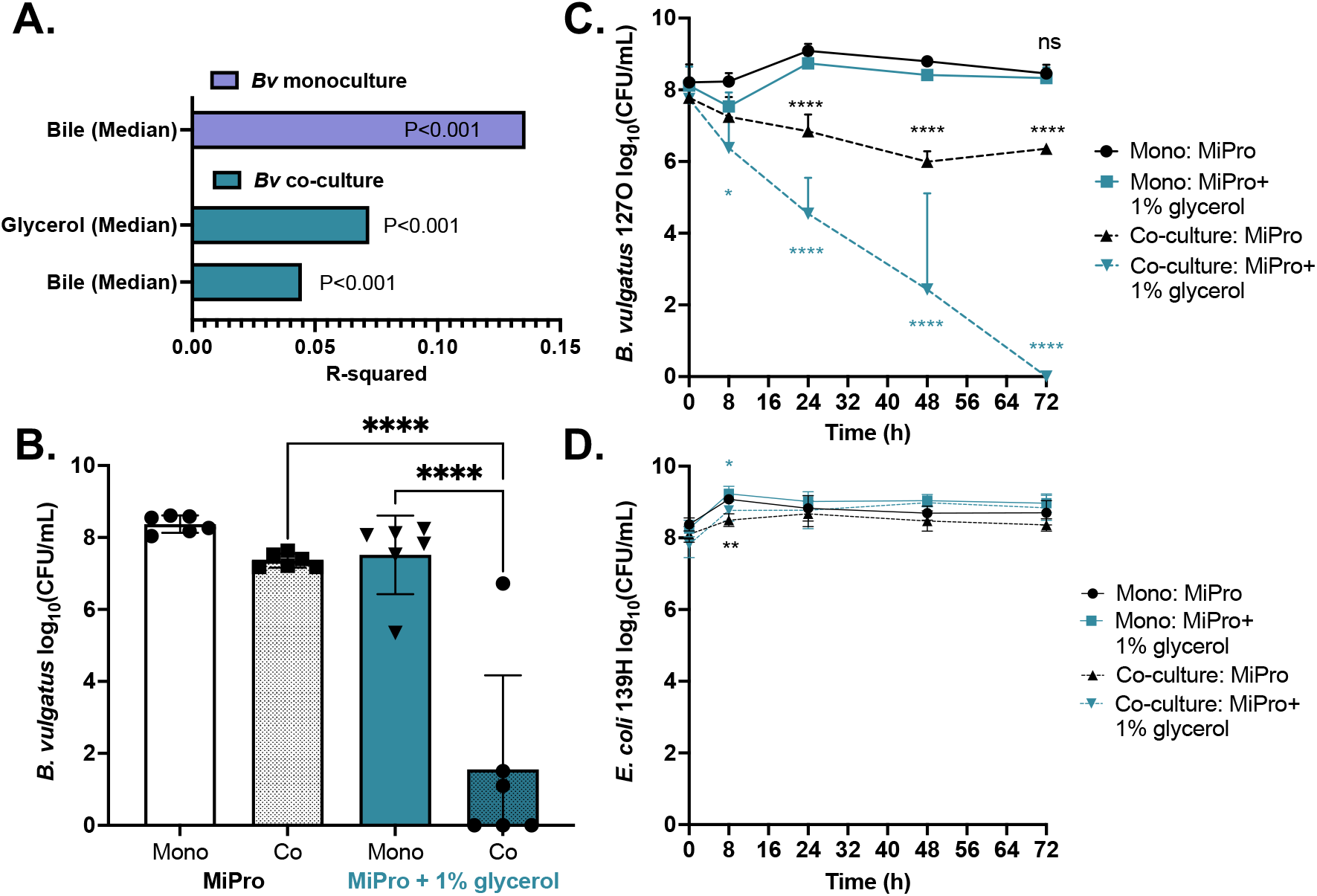
Linear regression identifies key components of CF gut physiology that contribute to *B. vulgatus* growth defects. **(A)** Significant features that contribute to loss of viability of *B. vulgatus* (*“Bv”*, strain n=6) in monoculture (purple, top) and in co-culture with *E. coli* (strain n=8; teal, bottom), compared to the MiPro growth condition. “Median” denotes the concentration of each feature to be at median-CF-MiPro levels. R-squared values explain the contribution to variance in the model of each feature. **(B)** Endpoint survival of *Bv* strains in monoculture (“Mono”) or co-culture with *E. coli* 139H (“Co”) for 72h growth in MiPro (black/white) or MiPro supplemented with 1% glycerol (teal). Each point denotes an individual *Bv* strain. Statistical analysis performed using ordinary one-way ANOVA with Tukey’s multiple comparisons. **(C)** Growth kinetics of one *Bv* strain, 127O, in monoculture (solid lines) or co-culture with *E. coli* 139H (dashed lines) in MiPro (black) or MiPro+1% glycerol (teal) over the course of 72h. Comparative *E. coli* 139H growth kinetics are displayed in **(D)**. Statistical analysis performed using 2-way ANOVA with Šídák’s multiple comparisons between co-culture and monoculture CFU/mL within the same growth medium (^*^, p < 0.05; ^**^, p < 0.01; ^***^, p < 0.005; ^****^, p < 0.001).

In all models, the tested features have relatively low contribution to overall variation (adjusted R^2^), ranging from 14.4 to 26.6%. This finding suggests that, individually, the CF-relevant features account for modest shifts in growth. However, in the context of the CF gut, these features are working in combination with each other, which likely compounds the observed growth defects. Thus, a more comprehensive approach is needed to properly address this question.

Bile’s negative impact on *B. vulgatus* survival was initially surprising, as *Bacteroides* species are generally considered bile-tolerant or bile-resistant [31]. In fact, the selective medium used to isolate *Bacteroides fragilis*, bile esculin agar, contains high concentrations of bile [32]. These microbes possess enzymes that can modify bile acid chemistry, either through deconjugation or conjugation with alternative amino acids in the case of bile salt hydrolases (BSHs) [33]. It is thought that these modifications detoxify primary conjugated bile acids to promote colonization [34], however other explanations focus on nutrition and energy usage of the liberated amino acids [35, 36]. Conversely, bile acid modifications may lead to more toxic metabolites [37], suggesting an array of putative outcomes attributed to microbiota metabolism of bile. A recent study reported an inhibitory effect of deoxycholic acid (DCA) on *Bacteroides* spp., including *B. vulgatus*, partially attributed to its intracellular accumulation [38]. The authors also note that inhibitory effects of bile acids are strain-specific, which agrees with our *B. vulgatus* monoculture growth data in median bile concentrations (**Supplemental Table 8**). The impacts of bile on *B. vulgatus* persist in co-culture with *E. coli*, suggesting bile as a microbial stressor regardless of competition.

We decided to pursue the role of glycerol (a surrogate of fat) on modulating *B. vulgatus* growth for two reasons: (1) to our knowledge, glycerol has not yet been implicated in the depletion of *Bacteroides* in the context of CF, and (2) glycerol only reduces *B. vulgatus* viability when *E. coli* is present, suggesting the role for microbial competition. First, we sought to quantify free glycerol in stool from CF (n=10) and nonCF (n=12) children to reaffirm its appropriate use as a fat surrogate. Free glycerol is not incorporated into triacylglycerols (TAGs) and is thus a direct measure of available glycerol in stool rather than what is hydrolyzed from TAGs in the processing of the fecal extracts for analysis. One mechanism by which free glycerol may accumulate in the CF intestine is due to a lack of absorption following liberation from TAGs by pancreatic or microbial lipases, however this point has not been explored. Using processing methods lacking saponification (e.g., strong base or acid) to avoid hydrolysis of glycerol from TAGs and ultra performance liquid chromatography/tandem mass spectrometry (UPLC–MS/MS), we successfully quantified free glycerol in stool samples from cwCF with healthy age-matched controls (**Supplemental Figure 2**). NonCF stool samples averaged 0.01675 μg/mg free glycerol and CF stool samples averaged 0.1515 μg/mg free glycerol (P=7.4e-05), signifying an 8-fold increase of compound in CF stool. Statistical analysis was performed using a mixed effect linear model to account for genotype, with age of donor and batch effects controlled for as the random variables.

Next, we investigated the dynamics of glycerol-mediated reduction of *B. vulgatus* viability in the presence of *E. coli*. At 72h, there is a significantly reduced *B. vulgatus* viability in co-culture with *E. coli* when glycerol is supplemented in MiPro pH 7 (**Figure 2B**). This depletion is observed in almost all strains of *B. vulgatus*, with 50% of strains exhibiting viability below detection. We confirmed that the low viability is independent of enhanced *E. coli* growth in glycerol (**Supplemental Figure 3**). We sought to track the death curve of one of these *B. vulgatus* strains, 127O, over the course of 72h co-culture with *E. coli* 139H (**Figure 2C**). Here, we observe 127O depletion as early as 8h, with enhanced killing at 24h compared to the MiPro pH 7 co-culture comparison. By 72h, 127O CFU/mL are below detection. This reduced viability is independent of enhanced *E. coli* 139H in glycerol over the time course (**Figure 2D**).

Interestingly, this interaction is exclusive to live cell cultures; cell-free supernatants from *E. coli* 139H monocultures or co-cultures (with *B. vulgatus* strain CFPLTA003-2B) do not recapitulate the reduction of *B. vulgatus* (**Supplemental Figure 4**). We tested anoxic monoculture supernatants from (1) *E. coli* 139H grown in both MiPro pH 7 and MiPro pH 7+1% glycerol for 72h and (2) *E. coli* 139H and *B. vulgatus* CFPLTA003-2B in MiPro or MiPro+1% glycerol for 72h, exposed *B. vulgatus* strains (n=5) and *E. coli* 139H in monoculture to diluted fractions of the supernatants and quantified growth via CFU/mL after 24h (**Supplemental Figure 4**). None of the tested supernatant conditions inhibited *B. vulgatus* in monoculture, compared to the control conditions lacking supernatant. The only exception observed was the supernatant from *E. coli-B*.*vulgatus* co-culture in MiPro slightly reduced *B. vulgatus* 127O viability in monoculture, but not to the same degree as observed with live cells. These results suggest that the reduction of *B. vulgatus* viability observed in the presence of glycerol and *E. coli* is likely not due to a stable secreted metabolite or toxin, but rather requires cell-to-cell contact. Alternatively, a secreted metabolite or toxin may be involved in this interaction, but this molecule is either unstable or it is transiently acting on *B. vulgatus*. Next, we aimed to explore the mechanism underlying the reduction of *B. vulgatus* viability by *E. coli* in the context of glycerol.

### Polyketide synthase (*pks*)-encoded colibactin is involved in modulating the *E. coli - B. vulgatus* interaction in the presence of glycerol

To address the mechanism of *E. coli-B. vulgatus* competition, we performed a genetic screen to identify *E. coli* mutants that do not reduce *B. vulgatus* viability in co-culture with *E. coli* 139H when grown in MiPro + 1% glycerol. To do so, *E. coli* 139H was conjugated as recipient with an *E. coli* strain donor containing a plasmid harboring the Tn*M* mariner transposon. The *E. coli* strain donor is auxotrophic for diaminopimelic acid (DAP), an amino acid precursor in the biosynthesis of lysine. Thus, we select for *E. coli* 139H mutants with transposon insertion by growth on lysogeny broth (LB) agar supplemented with gentamycin (selection) lacking DAP (counter-selection). A total of nearly 10,000 colonies were generated in this mutant library, representing 2x coverage of the 139H genome (**Supplemental Table 9**).

Individual insertion mutants were screened for their ability to grow similarly to the *E. coli* 139H WT parent in monoculture and co-culture with *B. vulgatus* 127O in MiPro + 1% glycerol after 72h anaerobic growth, but do not reduce the viability of *B. vulgatus* 127O growth in co-culture relative to its monoculture control. A visual schematic of the transposon screen procedure is outlined in **Supplemental Figure 5**. Candidates that demonstrated this phenotype with *B. vulgatus* 127O in co-culture, were denoted by “full”, “partial” or “minimal” restored viability relative to *B. vulgatus* 127O monoculture controls, were selected for secondary and tertiary validation screens.

We obtained 549 primary candidates (5.56% of screened colonies), with a range of rescue capabilities: 87 (0.88%) of mutants restored *B. vulgatus* 127O viability fully relative to monoculture, 173 (1.75%) mutants restored *B. vulgatus* 127O viability partially, and 289 (2.93%) mutants restored *B. vulgatus* 127O viability minimally (**Supplemental Table 9**). Only 51/549 candidates were validated in the secondary screen to restore *B. vulgatus* 127O viability fully. For the primary “full restoration” candidates (n=87) that displayed only partial restoration of viability in the secondary screen, we denoted these as secondary “partial restoration” candidates, of which there were 24.

The location of transposon insertion in the candidate strains was determined via arbitrary primed PCR and Sanger sequencing of flanking regions, as described in the Materials and Methods. From this screen, we identified candidates that were categorized into genes related to: metabolism, stress response, biofilm formation and its regulation, phage, rRNA and pathogenicity (**Supplemental Table 10-11**). Of note, one full restoration candidate (*dhaR*::Tn*M*) and three partial restoration candidates (*dhaK*::Tn*M, dhaI*::Tn*M, gldA*::Tn*M*) are related to glycerol metabolism, serving as sound positive controls confirming the validity of the screen and the potential for glycerol metabolic by-products (i.e., methylglyoxal) as potential toxic mediators in the *E. coli-B*.*vulgatus* interaction. Other metabolic genes with transposon insertions include *actP*, an acetate symporter, *malE*, a maltose binding protein, and *alsK*, an allose kinase. These candidates likely suggest the importance of nutritional competition of these substrates in the microbial interaction. Other strains with full restoration phenotypes had mutations in genes involved in stress responses to either oxidative stress (*sodA*::Tn*M*) or misfolded proteins (a Tn*M* in the *clpQY* promoter), and biofilm maintenance (*pdeH*::Tn*M*) or regulation (*rsmE*::Tn*M*), suggestive of other mechanisms of competition involving stress tolerance by means of detoxification or biofilm formation.

However, the most striking result from this screen was that 51% of the validated “full restoration” candidates involved the biosynthesis of colibactin, a genotoxin associated with increased colorectal cancer risk [39-43]. The *pks*, or polyketide synthase, gene cluster is a 54 kb genetic island that consists of 19 *clb* genes encoding various non-ribosomal peptide synthetase (NRPS), polyketide synthetase (PKS), hybrid NRPS-PKS and other biosynthetic enzymes as well as accessory proteins [25, 41]. Colibactin production is largely regulated by the first two genes in the cluster, *clbA* and *clbR*, and sets of genes in the cluster are transcribed independently. Specifically, there are four polycistronic elements (1: *clbA-clbR*, 2: *clbCDEFG*, 3: *clbIJKLMN*, and 4: *clbOPQ*) and three monocistronic elements (*clbB, clbH, clbS*). When expressed, the biosynthetic scaffold coordinates production of the inactive biosynthetic intermediate precolibactin, which contains two *N-*myristoyl- d-Asn motifs. After export of precolibactin from the cytoplasm, the biosynthetic enzyme ClbP, a periplasmic transmembrane peptidase, cleaves the prodrug motifs, triggering production of the active colibactin molecule, characterized by two electrophilic cyclopropane warheads [44-46]. These warheads alkylate DNA at adenine-rich motifs (5’-AAWWTT-3’, where W = A or T), causing DNA interstrand cross-linking and activation of DNA repair machinery [47-49].

The 26 independent transposon insertions spanned across five genes in the 54 kb *clb/pks* island, with single hits in *clbD* and *clbN*, and multiple hits in *clbB, clbJ* and *clbK* (**Figure 3A, Supplemental Table 11**). 20 of the 26 transposon insertions are oriented in the 3’ to 5’ direction, or antisense, to the coding region. In addition to disrupting the function of the gene in which the transposon is inserted, the outward facing P_*tac*_ promoter may be modulating the expression of the gene upstream of the insertion in these cases. In two independent candidates, the transposon is detected in both *clbJ* and *clbK*, denoted as “linked” with a bracket (**Figure 3A**). In these strains, it appears that there are two transposon insertions: one in *clbJ* and one in *clbK*. Furthermore, two “partial restoration” candidates (*htpG*::Tn*M*) are molecular chaperones that aid in colibactin synthesis (**Supplemental Table 10**) [50], yet are not transcribed from the *pks* genetic island, further enforcing the role of colibactin synthesis in this phenotype.

**Figure 3.**
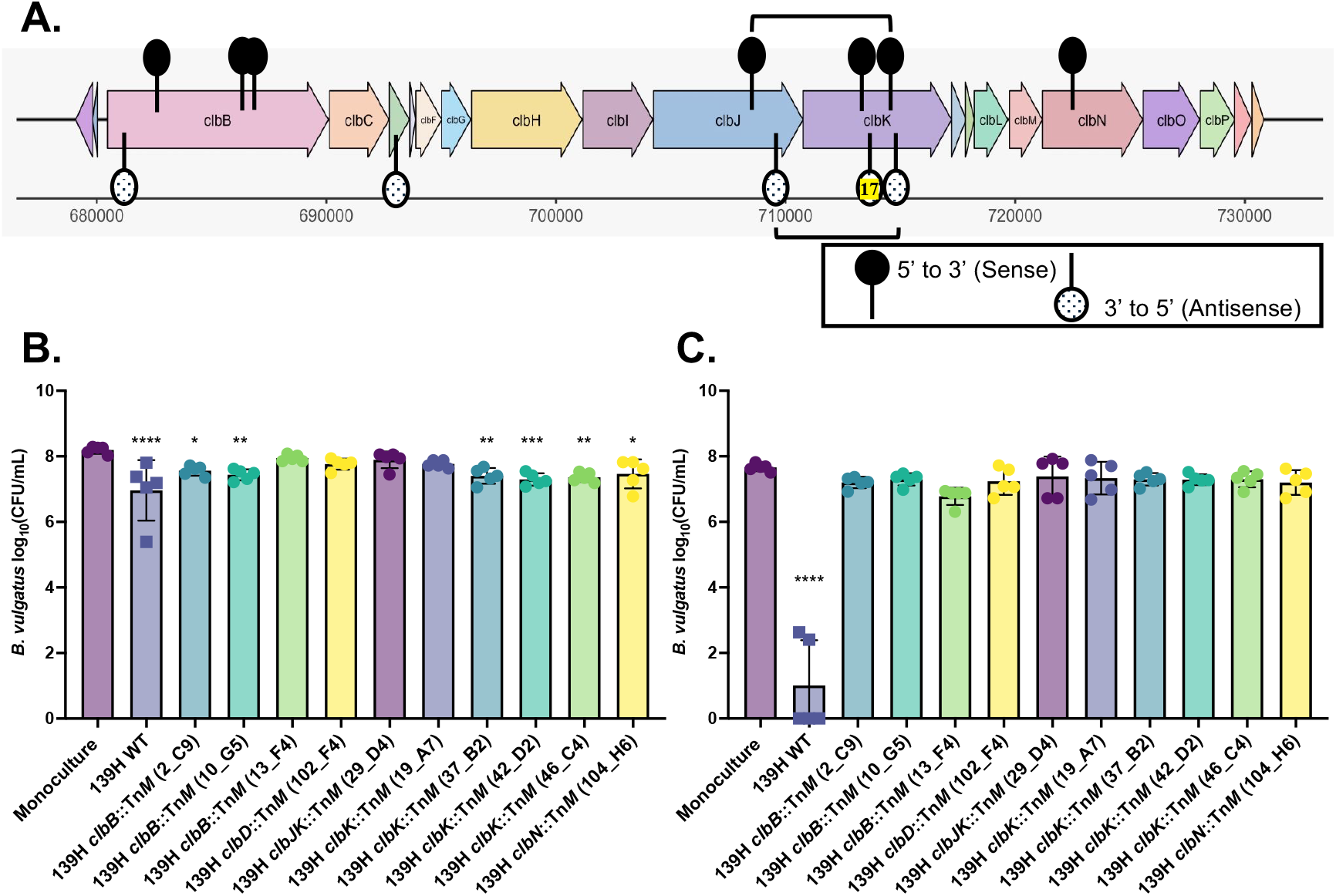
Transposon mutagenesis identified colibactin as necessary in *B. vulgatus* viability reduction by *E. coli* in the presence of glycerol. **(A)** Schematic of the *pks/clb* genetic island, spanning 54 kb and containing genes *clbA – clbS*. Black lollipops denote transposon insertions in the 5’ to 3’ direction, inverted white and black checkered lollipops denote transposon insertions in the 3’ to 5’ direction. There were 17 independent antisense hits to *clbK*, as represented by the highlighted ‘17’. Two independent candidates have transposon insertions detected in both *clbJ* and *clbK*, described as “linked” and denoted with the bracket. **(B-C)** *B. vulgatus* clinical isolates (n=5) grown in monoculture (first bar, purple) and co-culture with *E. coli* 139H WT (second bar, navy) and representative *E. coli clb*::Tn*M* mutants for 72h in **(B)** MiPro pH 7 and **(C)** MiPro pH 7 + 1% glycerol. Statistical analysis performed using one-way ANOVA with Dunnett’s multiple comparisons between each co-culture condition with the monoculture control (^*^, p < 0.05; ^**^, p < 0.01; ^***^, p < 0.005; ^****^, p < 0.001).

First, we aimed to confirm that the transposon mutants grew to the same extent as *E. coli* 139H WT. To do so, we selected at least one representative mutant from each gene that was hit and cultured these strains in monoculture for 72h anaerobically in MiPro pH 7 and MiPro pH 7 + 1% glycerol. There are no observed growth advantages in glycerol-supplemented medium across all strains, including WT (**Supplemental Figure 6A**). However, one of the *clbJK*::Tn*M* “linked” candidates (29_D4) exhibits a slight but significant growth defect at the CFU/mL level compared to WT in MiPro + 1% glycerol. This observation may contribute to *B. vulgatus* 127O’s ability to not show a reduction in co-culture with this mutant.

Next, we performed a tertiary validation screen with the representative *clb*::Tn*M* hits to confirm a full restoration of *B. vulgatus* 127O viability by determining CFU/mL for both microbes. In addition to *B. vulgatus* 127O, we tested 4 other strains of *B. vulgatus* to address whether this restoration of viability is applicable across strains. First, we co-cultured each *B. vulgatus* strain with *E. coli* 139H WT and with each *E. coli* Tn*M* mutant in MiPro pH 7 for 72h and compared co-culture growth to the monoculture control (**Figure 3B**). The WT strain significantly inhibits *B. vulgatus* strains in co-culture by ∼1000 CFU/mL, while some of the transposon mutants showed restored *B. vulgatus* viability, with the exception of mutants *clbB*::Tn*M (*2_C9 and 10_G5), *clbK*::Tn*M* (37_B2 and 42_D2, 46_C4) and *clbN*::Tn*M* (104_H6). These transposon mutants significantly reduce *B. vulgatus* viability in co-culture, relative to monoculture controls, however to a lesser degree than *E. coli* 139H WT. We repeated these co-culture assays in MiPro pH 7 + 1% glycerol and observed complete restoration of viability for all *B. vulgatus* strains across all tested transposon mutants (**Figure 3C**). Thus, the restoration of viability of *B. vulgatus* is more consistent in glycerol (e.g., the conditions in which the mutant screen was performed) and translatable across all tested *B. vulgatus* strains. Despite one transposon mutant (*clbJK*::Tn*M* 29_D4) with a slight but significant growth defect compared to WT, all tested *E. coli* 139H mutants grow similarly to WT in co-culture with *B. vulgatus* strains in both MiPro and MiPro supplemented with glycerol (**Supplemental Figure 6B-C**), suggesting that changes in endpoint *E. coli* CFU/mL are not responsible for driving *B. vulgatus* rescue.

Finally, we tested the growth kinetics of each representative transposon insertion against *E. coli* 139H WT to account for potential differences in log-phase growth. Here, we inoculated *E. coli* 139H WT and each representative transposon mutant in optically-clear supplemented brain heart infusion broth (BHIS) buffered to pH 7 and BHIS pH 7 + 1% glycerol at OD_600_ = 0.01. Using a spectrophotometer in an anaerobic chamber at 37 °C, we quantified OD_600_ of the strains in each condition every 3 minutes for 8h. All strains grow similarly in BHIS pH 7, with no changes compared to the WT strain (**Supplemental Figure 7A**). In glycerol-supplemented medium, all transposon mutants, except for *clbK*::Tn*M* (37_B2), grew similarly to WT; the *clbK*::Tn*M* mutant has enhanced growth in late log-phase (2.9h – 3.4h) compared to WT, however endpoint OD_600_ between the two strains are similar (**Supplemental Figure 7B**). These results suggest that the restoration of *B. vulgatus* viability by these transposon mutants is independent of observed growth defects in the mutant strains compared to growth with *E. coli* 139H WT.

### Glycerol and microbial competition modulate colibactin production in vitro

We next sought to investigate the role of glycerol and microbial competition in the production of colibactin to address whether this molecule is induced or repressed by these factors. Literature suggests that specific environmental and metabolic factors modulate expression of the *pks* island, including iron concentration [51, 52], oligosaccharides (i.e., glucose and inulin) [53, 54], short-chain fatty acids [55, 56], oxygen [57], serine [58], spermidine and inflammation through unknown mechanisms [41]. These *pks*-enhancing factors can be attenuated through the addition of inhibitory factors (i.e., ferrous sulfate in the case of iron [54]), suggesting that metabolic conditions and colibactin production regulation are linked. To the best of our knowledge, glycerol has yet to be investigated as a potential modulator of colibactin regulation.

To test the hypothesis that glycerol coupled with microbial competition induces colibactin production, we performed ultra-performance liquid chromatography tandem mass spectrometry (UPLC–MS/MS) to quantify the prodrug scaffold that is hydrolyzed by the ClbP periplasmic peptidase during colibactin activation. The prodrug scaffold, *N*-myristoyl-d-Asn, is stable under aerobic conditions compared to the final colibactin molecule [57], thus rendering it useful for quantification studies [59].

First, we examined the difference in prodrug production across several *E. coli* strains: Nissle (*pks+*), 143D (*pks-*), 139H (*pks+*), 139H *clbN*::Tn*M* (*pks*-mutant^*^), NC101 (*pks+*) and NC101 Δ*clbP* (*pks*-mutant^*^). Monocultures of each isolate were grown anaerobically in MiPro and MiPro supplemented with 1% glycerol for 24h. Cell count was quantified via CFU/mL and whole-culture prodrug concentrations were normalized to cell count. Overall, the *E. coli pks+* strains (e.g., Nissle, 139H and NC101) produce moderate amounts of prodrug in MiPro (∼11-18 nM/CFU mL^-1^; **Figure 4A**), and this production is significantly increased in the presence of glycerol (∼72-89 nM/CFU mL^-1^; **Figure 4A**), with the exception of Nissle (21 nM/CFU mL^-1^). Notably, the *pks* mutants (e.g., 139H *clbN*::Tn*M* and NC101 Δ*clbP*) produce prodrug levels below the limit of detection, similar to levels observed in the *pks-* strain 143D. Taken together, these results suggest that glycerol induces the production of colibactin (except for the Nissle strain), as quantified by measuring levels of *N*-myristoyl-d-Asn. Additionally, these findings confirm the elimination of colibactin production in both a *clbN* transposon mutant and a *clbP* null mutant.

**Figure 4.**
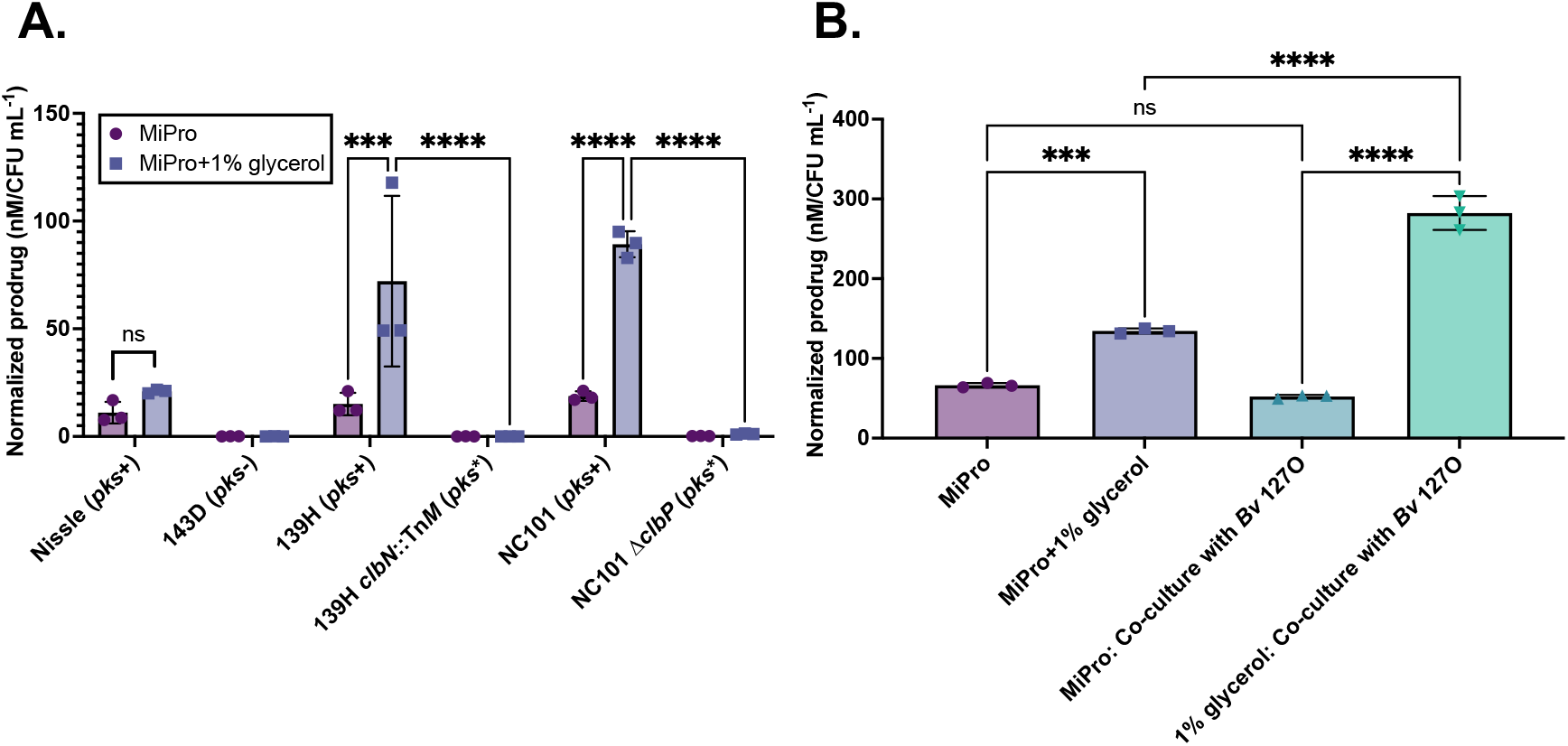
Glycerol and microbial competition modulate colibactin prodrug production in vitro. **(A)** *E. coli* strains (n=6) were grown anaerobically in MiPro pH 7 (circles) and MiPro pH 7+1% glycerol (squares) for 24h. Following incubation, cell count was quantified via CFU/mL plating on LB agar, and levels of *N*-myristoyl-d-Asn, the colibactin-derived prodrug motif (labeled “prodrug”) were quantified in whole cultures using LC-MS. Prodrug concentrations were normalized to cell count (nM/CFU mL^-1^) across all conditions. Statistical analysis performed using 2-way ANOVA with Šídák’s multiple comparisons (^***^, p<0.005; ^****^, p<0.001). **(B)** *E. coli* 139H was grown anaerobically in MiPro pH 7 and MiPro pH 7+1% glycerol with and without co-culture with *B. vulgatus* 127O for 72h. Following incubation, cell count was quantified via CFU/mL plating on LB agar for *E. coli* and blood sheep agar+100 µg/mL gentamicin for *B. vulgatus*. Levels of *N*-myristoyl-d-Asn, the colibactin-derived prodrug motif (labeled “prodrug”) were quantified in whole cultures using LC-MS. Prodrug concentrations were normalized to *E. coli* cell count (nM/CFU mL^-1^) across all conditions. Statistical analysis performed using one-way ANOVA with Šídák’s multiple comparisons (^***^, p<0.005; ^****^, p<0.001).

Next, we investigated the role of microbial competition on colibactin production. *E. coli* 139H was grown in monoculture and co-culture with *B. vulgatus* 127O anaerobically in MiPro and MiPro supplemented with 1% glycerol for 72h. Cell count was quantified via CFU/mL to confirm a reduction in *B. vulgatus* viability, and whole-culture prodrug concentrations were normalized to cell count of *E. coli*. Similar to what was observed in 24h monocultures, glycerol induces production of prodrug in 72h cultures compared to MiPro alone (P<0.0001; **Figure 4B**). Interestingly, co-culture with *B. vulgatus* in MiPro does not change prodrug concentrations relative to the MiPro monoculture control (P=0.43). However, prodrug concentrations in co-cultures in 1% glycerol are significantly increased compared to both the glycerol monoculture control (P<0.0001) and MiPro co-culture (P<0.0001; **Figure 4B**), suggesting that both glycerol and microbial competition are required to maximize colibactin biosynthesis.

### *pks+ E. coli* strains are found in both nonCF and CF infants and correlate with the depletion of *B. vulgatus*

Next, we explored the prevalence of *pks*-positive strains of *E. coli* in the context of the CF gut microbiome with the hypothesis that increased detection or isolation of colibactin-encoding strains of *E. coli* may lend increased risk of altered microbial interactions in CF gut microbiomes. First, we aimed to characterize our collection of CF and nonCF *E. coli* clinical isolates for the colibactin biosynthetic cluster. To date, we have 22 isolates (CF: 6, nonCF: 15, lab: 1) originating from children’s stool or adult colonoscopy samples that have been whole genome sequenced. For the CF collection, 4/6 strains were isolated from children’s stool (n=3 children), while 2/6 strains were isolated from a single adult CF colonoscopy sample (GIEHZ100). For the nonCF collection, 15/15 strains were isolated from a single adult nonCF colonoscopy sample (GIEHV103). Interestingly, the *E. coli* lab strain (Nissle 1917), which is a recognized probiotic strain [60], contains the complete 19-gene *pks* island, including an annotated *clbS/dfsB* family gene that may or may not have related or redundant function to the immunity protein *clbS*. Given this homology, we included this strain in this study. We used the protein sequences from the complete Nissle *pks* island as the reference in our comparative genomics study.

Using BLAST, we compared the Nissle Clb protein sequence to query the whole genome of each clinical isolate as the subject sequence. We determined which clinical isolates possessed homologous protein sequences to each Clb protein in the cluster, the percent identity, e-value and other quality parameters compared to the Nissle reference, as well as where in the genome the island was located. We filtered the sequences by e-value (≤ 1E-10) and percent identity (≥ 50%); the clinical isolate sequences that did not meet this filtering threshold were marked as absent for that protein (**Supplemental Table 12**). Next, we generated a heat map to display sequence similarity of the colibactin biosynthetic cluster across isolates (**Supplemental Figure 8**). The results from this comparative genomics analysis show that two CF and one nonCF stool *E. coli* isolates are devoid of any Clb proteins (138G,143D and GP0236). One CF stool isolate (139G) and two CF colonoscopy isolates (GP0165 and GP0192) possess various proteins in the cluster with relatively high sequence similarity to the Nissle sequence. However, these partial sequences all lack *clbA* and *clbR*, the two regulators of the cluster. The genes detected in GP0165 and GP0192 have 100% sequence similarity to each other, supporting the clonal nature of strains isolated longitudinally in the GI tract in a single person. Interestingly, only one CF isolate contains the full *pks* island (*E. coli* 139H). This strain and *E. coli* 139G were isolated from stool from the same child with CF within a three-month span, suggesting acquisition or adaptation of colibactin production may arise in CF childhood. In the nonCF strain collection, 14/15 strains contain the full *pks* island. Only one strain, GP0236, lacks all proteins in the cluster. This strain was isolated from the ascending colon aspirate of the nonCF colonoscopy donor, along with *pks+* strains *E. coli* GP0234 and GP0247. While most isolates from this donor appear clonal at the *pks* level, this result suggests that there may be populations of *E. coli* with various degrees of colibactin conservation within one GI tract.

These results reveal that colibactin-producing strains of *E. coli* can be isolated from gut microbiome samples of both CF and nonCF genotypes. However, the sample size of patients from which these strains were isolated is low (CF: 5, nonCF: 1). Additionally, we isolated these strains via culture-dependent methods, which may introduce culture-based biases. Lastly, the demographics of the two genotype cohorts are very different, with respect to age; the CF cohort is 80% children, while the nonCF cohort is 100% adult. Because there is evidence that *E. coli* may acquire the colibactin biosynthetic cluster via horizontal gene transfer [61], age and *E. coli* abundance likely play a role in detection. Thus, to better examine the role of colibactin on microbial competition, we performed in vitro competition assays with *B. vulgatus* and the majority of our *E. coli* clinical isolate collection with the hypothesis that *pks+ E. coli* strains will reduce *B. vulgatus* viability more than *E. coli* strains lacking these genes. Results from the co-culture assays suggest that, in MiPro, the viability of *B. vulgatus* strains are modestly reduced by *E. coli* strains that possess the full suite of *clb* genes (+), represented by the strains labeled with dark purple bars (*E. coli* 139H, GP0281, GP0247, GP0306, GP0234, GP320, Nissle; **Figure 5A**). *E. coli clb+* strains that reduce *B. vulgatus* viability in co-culture relative to monoculture controls in MiPro are *E. coli* GP0234 and GP0247. Other *E. coli* strains that contain only some of the *clb* genes (/), represented by strains labeled with light purple bars, or none of the *clb* genes (-; white bars) do not reduce *B. vulgatus* viability when co-cultured in MiPro. In MiPro + 1% glycerol, all *E. coli clb+* strains significantly reduce *B. vulgatus* viability, relative to its monoculture growth (**Figure 5B**). In contrast, there is no observed reduced viability by *E. coli clb(/)* or *clb(-)* strains. This observation is independent of the source of the isolate and not associated with changes in *E. coli* growth (**Supplemental Figure 9A-B**). The one exception is observed phenotype is *E. coli* Nissle, which is *clb+*. This strain does not inhibit *B. vulgatus* under either growth condition. This observation is likely explained by the reduced concentration of prodrug produced by Nissle in glycerol-supplemented medium (**Figure 4A**). In addition to the reduction in prodrug production by this strain, Nissle may also produce metabolites, substrates or other factors that support *B. vulgatus* growth in co-culture and ameliorate microbial competition with resident microbes. This phenomenon has been explored in numerous studies which generally support the use of Nissle as a probiotic strain, despite its *pks+* status [62, 63].

**Figure 5.**
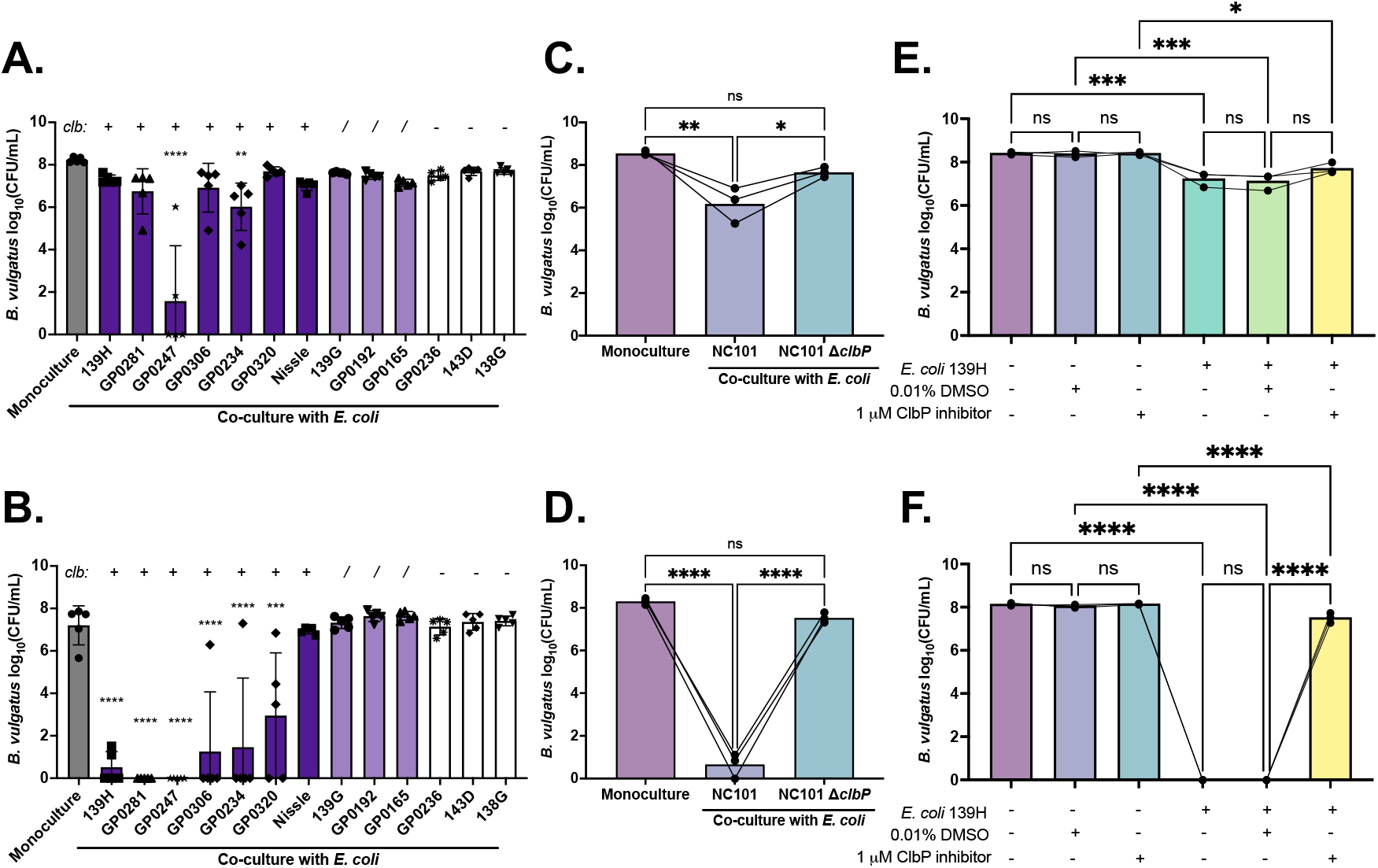
The *pks* gene cluster in *E. coli* correlates with reduced *B. vulgatus* viability in the presence of glycerol. **(A-B)** *B. vulgatus* strains (n=5) grown anaerobically in monoculture (first grey bar) and co-culture with 13 individual *E. coli* isolates in **(A)** MiPro pH 7 and **(B)** MiPro + 1% glycerol for 72h. *E. coli* strains with a complete *pks/clb* gene cluster are colored in dark purple (+), partial *pks/clb* gene cluster in light purple (/), and no *pks/clb* genes in black/white (-). Statistical analysis performed using one-way ANOVA with Dunnett’s multiple comparisons between co-culture CFU/mL to monoculture CFU/mL (^*^, p < 0.05; ^**^, p < 0.01; ^***^, p < 0.005; ^****^, p < 0.001). **(C-D)** *B. vulgatus* strains (n=3) grown anaerobically in monoculture (first purple bar) and co-culture with *E. coli* NC101 (second light purple bar) and NC101 Δ*clbP* (third blue bar) in **(C)** MiPro pH 7 and **(D)** MiPro + 1% glycerol for 72h. The average log_10_(CFU/mL) of 3 biological replicates of each *B. vulgatus* strain are connected by a line. Statistical analysis was performed using one-way ANOVA with Tukey’s multiple comparisons (^*^, p<0.05; ^**^, p<0.01; ^****^, p<0.001). **(E-F)** *B. vulgatus* strains (n=3) grown anaerobically in monoculture (1^st^, 3^rd^ and 5^th^ bars; “*E. coli* 139H –”) in **(E)** MiPro pH 7 or **(F)** MiPro pH 7+1% glycerol supplemented with 0.01% DMSO (3^rd^ and 4^th^ bars) or 1 µM ClbP inhibitor (5^th^ and 6^th^ bars). The table below represents presence (+) or absence (-) of co-culture with *E. coli* 139H, supplementation with 0.01% DMSO vehicle or 1 µM ClbP inhibitor. The average log_10_(CFU/mL) of 3 biological replicates of each *B. vulgatus* strain are connected by a line. Statistical analysis was performed using one-way ANOVA with Tukey’s multiple comparisons (^*^, p<0.05; ^***^, p<0.005; ^****^, p<0.001).

Next, we aimed to test the necessity of colibactin production in the observed interaction with *B. vulgatus*. To do this, we applied both genetic and chemical approaches. First, we obtained an *E. coli* strain lacking *clbP*, which encodes the periplasmic peptidase responsible for hydrolyzing the prodrug and activating colibactin. We co-cultured three independent *B. vulgatus* strains with the *clbP* mutant and its parent strain, NC101, in MiPro pH 7 (**Figure 5C**) and MiPro pH 7+1% glycerol (**Figure 5D**). Results from these co-cultures suggest that NC101, a *pks+* strain of *E. coli*, reduces viability in all tested isolates of *B. vulgatus* in MiPro, and this reduction is exacerbated in glycerol-supplemented medium. Furthermore, when *clbP* is absent, this reduction in viability is rescued similar to monoculture controls in both growth conditions. *E. coli* cell counts remain largely unchanged compared to its monoculture growth control (**Supplemental Figure 9C-D**), supporting the lack of association with *E. coli* growth.

We also tested the role of colibactin through small molecule inhibition of ClbP [59]. This previously reported inhibitor is derived from a boronate ester precursor, (*S*)-*N*-(3-amino-3-oxo-1-(4,4,5,5-tetramethyl-1,3,2-dioxaborolan-2-yl)propyl)-4-phenylbutanamide, which spontaneously hydrolyzes in water to yield the active boronic acid inhibitor, (*S*)-(3-amino-3-oxo-1-(4-phenylbutanamido)propyl)boronic acid. This compound binds with high potency and selectivity to ClbP and mimics the hydrogen-bonding interactions of precolibactin, thus disrupting the hydrolysis of this biosynthetic intermediate [59]. When *E. coli* 139H is incubated with 1 µM of the ClbP inhibitor for 72h, prodrug concentrations are significantly reduced both in the presence and absence of glycerol, compared to the DMSO vehicle control (**Supplemental Figure 10**), confirming the potency of the compound. Furthermore, when *E. coli* 139H is grown in anaerobic co-culture with three independent *B. vulgatus* strains in MiPro pH 7 supplemented with either DMSO vehicle control or 1 µM of the ClbP inhibitor for 72h, there is a modest yet nonsignificant rescue of *B. vulgatus* viability in the presence of the inhibitor (**Figure 5E**). Importantly, in co-cultures with 1% glycerol, the ClbP inhibitor significantly restores *B. vulgatus* viability nearly to monoculture levels (**Figure 5F**). Again, this restoration in viability is independent of changes in *E. coli* growth (**Supplemental Figure 9E-F**). Taken together, these findings support the sufficiency of colibactin production in the reduction in *B. vulgatus* viability in our system.

### Analysis of CF stool metagenomes suggest increasing trends in prevalence and abundance of *clb* genes

To begin to address the discrepancy in cohort demographics from which the *E. coli* strains above were isolated (e.g., difference in ages and sample type between CF and nonCF cohort), we sought to investigate the prevalence and abundance of colibactin-related genes in stool samples from children with and without CF. This analysis aimed to eliminate culture biases attributed to the strain isolation process and focus on one age demographic: children under the age of three years old. To do so, we utilized publicly-available stool metagenomic data of CF and nonCF children [6]. Additionally, we extracted DNA from CF and nonCF stool samples (n=18 for each genotype) and sent this DNA for metagenomic sequencing. In all, our dataset includes 208 CF and 2,560 nonCF stool metagenomes. In terms of analysis, human reads were filtered and microbial taxonomy was assigned. Differential relative abundance of each taxon was calculated between genotypes. Microbial gene families and pathways were also assigned using the UniRef90 database. Statistical analyses were performed using Wilcox rank-sum test to account for genotype.

We first were interested in the prevalence of detectable *clb* genes in stool metagenomes. We focused on *clbB* due to the frequency at which this gene was targeted by transposon candidates and the conservation of this gene in *pks+ E. coli* clinical isolates. We observed that 372/2,560 (14.5%) of nonCF stool metagenomes had detectable levels of *clbB*, compared to 60/208 (28.8%) of CF stool metagenomes (**Figure 6A**), suggesting that the percent of stool metagenomes with detectable levels of colibactin-related genes is higher in CF children than nonCF children. To our knowledge, this is the first study to identify an association between colibactin production and the CF genotype [26, 27]. Because our analysis is focused on children under the age of three years, these results may point to the application of *clb* gene detection as a biomarker for CRC in young cohorts.

**Figure 6.**
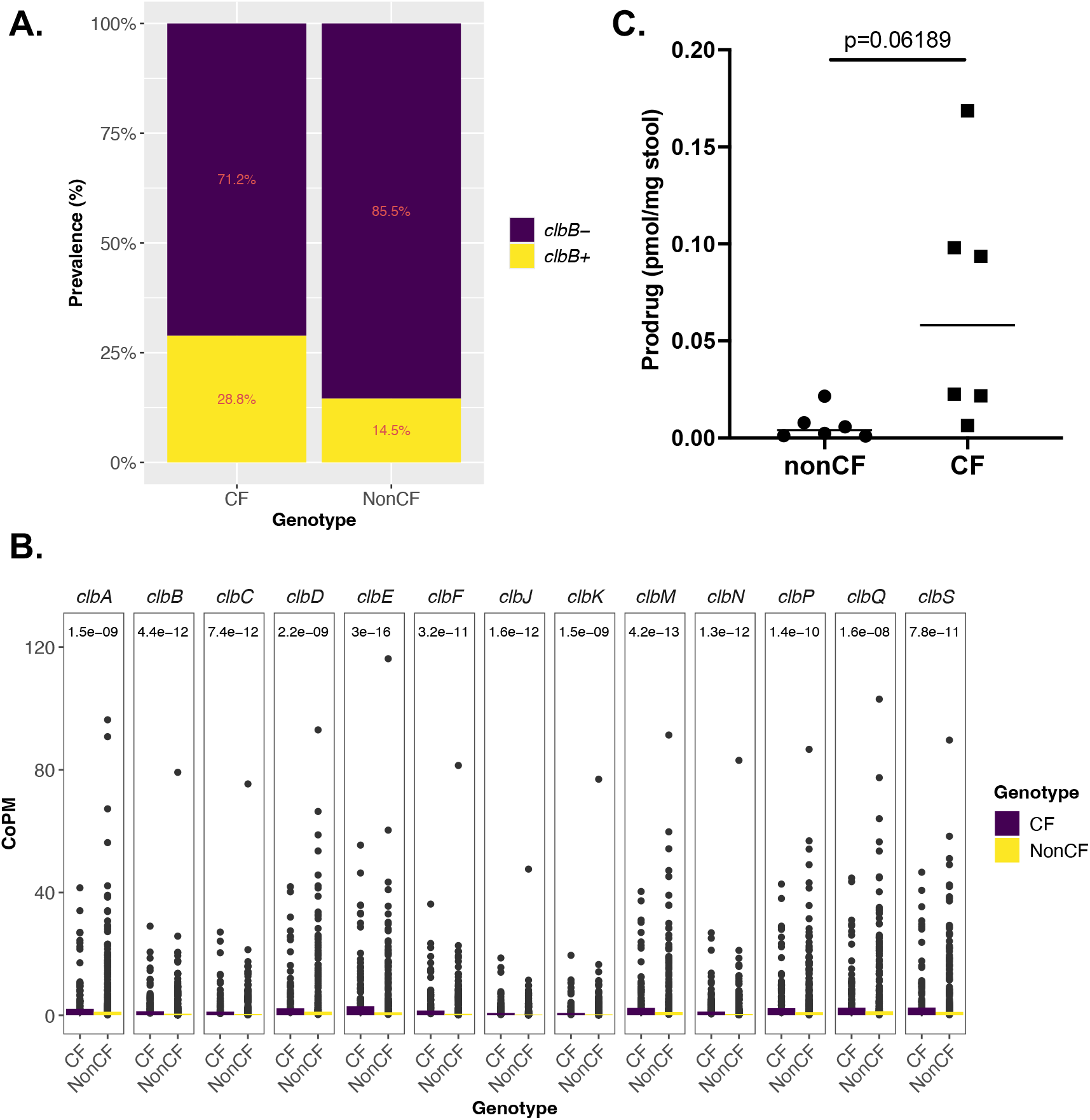
Markers of increased colibactin production are evident in CF stool. **(A)** Prevalence (%) of stool metagenomes with detectable *clbB* gene sequences (yellow, *clbB+*) or not (purple, *clbB-*) in CF and nonCF children. **(B)** Abundance for the majority of gene families in the *clb* island (*clbA, clbB, clbC, clbD, clbE, clbF, clbJ, clbK, clbM, clbN, clbP, clbQ, clbS*) not assigned to a particular bacterial species (“unclassified”) are first normalized for gene length with reads per kilobase and then normalized for library depth, as depicted by copies per million (CoPM). Each point represents the CoPM of a given gene in one sample. Mean gene counts are displayed for CF (purple) and nonCF (yellow) stool metagenomes in the filled bars. The mean gene count is higher in CF for each gene. Statistical analysis performed using Wilcox rank-sum test to account for genotype; p-values are displayed at the top of each gene facet. **(C)** The colibactin prodrug was quantified from raw stool collected from nonCF (n=6) and CF children (n=6). Concentrations were normalized to the weight of each stool sample (pmol/mg stool). Statistical analysis performed using mixed effect linear modeling with batch as the random variable, and genotype as the fixed variable.

Next, we aimed to quantify the gene abundance for the majority of *clb* genes in CF and nonCF stool metagenomes. Gene abundance values indicate frequency of a gene in a population. Our approach provides insight into the potential functional capabilities of colibactin production in these two microbiomes. Gene abundance for *clbA, clbB, clbC, clbD, clbE, clbJ, clbK, clbM, clbN, clbP, clbQ, clbS*, as displayed as mean copies per million (CoPM) reads [64], are significantly higher in CF stool metagenomes compared to those of nonCF (**Figure 6B**). Because there is also an enrichment of *E. coli* in the CF gut, we plotted *clbB* gene abundance against the relative abundance of *E. coli* and found a significant correlation (Pearson R^2^=0.44, P=0; **Supplemental Figure 11A**). Taken together, these results suggest that CF gut microbiomes possess higher levels of colibactin-associated genes, which are directly correlated with the relative abundance of *E. coli*.

Other species of Enterobacteriaceae, such as *Klebsiella pneumoniae*, are capable of synthesizing colibactin and contribute to gene counts in our dataset; however, the abundance of *clbB* gene counts does not correlate with the relative abundance of *K. pneumoniae* (Pearson R^2^= -0.04, P=0.545; **Supplemental Figure 11B**). Furthermore, because of our in vitro observations, we examined the correlation between *clbB* gene counts and *B. vulgatus* relative abundance. Similarly to *Klebsiella, clbB* gene counts do not correlate with levels of *B. vulgatus* in CF stool metagenomes (Pearson R^2^=0.01, P=0.835; **Supplemental Figure 11C**).

In nonCF stool metagenomes, a significant correlation between *clbB* and *E. coli* exists (Pearson R^2^=0.26, P=0; **Supplemental Figure 11D**), although to a lesser extent than that observed in CF stool metagenomes. These results may suggest contribution by other colibactin-producing bacteria, however we observed no correlation between *clbB* and *K. pneumoniae* (Pearson R^2^=0, P=0.897; **Supplemental Figure 11E**). Thus, these findings likely reflect differences in *E. coli* relative abundance between the two genotypes and may suggest an enhanced role for *E. coli-*mediated colibactin production in CF and its potential as a biomarker in this disease cohort. Interestingly, in nonCF stool metagenomes, there is a weak yet significant anticorrelation between *clbB* gene counts and *B. vulgatus* (Pearson R^2^= -0.06, P=0.03; **Supplemental Figure 11F**), lending support to the in vitro observation that *B. vulgatus* viability is reduced by *E. coli pks+* in when fat levels are not high.

Finally, we investigated the concentration of *N-*myristoyl-d*-*Asn, the colibactin prodrug motif, in CF and nonCF stool samples (n=6 each genotype). Here, we homogenized raw stool and quantified the prodrug motif using UPLC–MS/MS, analogous to quantification of the prodrug in *E. coli* cultures. The concentration was normalized to the weight of stool (nM/mg stool). Statistical analysis was performed using a mixed effect linear model to account for genotype (fixed variable) and batch effects (random variable). Here, we observe a trending increase in colibactin prodrug in CF stool compared to nonCF stool (P=0.062; **Figure 6C**). Taken together, these results present an enrichment of colibactin-encoding genes and a trending increase in the colibactin prodrug in CF stool compared to nonCF stool. Considering the association of colibactin with inflammation and the development of CRC, quantification of these markers in stool may be used for early detection of pathology in young CF cohorts.

## Discussion

In this study, we report that *Bacteroides*, specifically *B. vulgatus*, is sensitive to both CF-relevant in vitro growth conditions and microbial competition with *E. coli*. Under baseline conditions in a medium used to resemble the nutrients of the healthy intestine (MiPro) [28], strains of *B. vulgatus* show reduced viability starting at 8h and this reduction persists through 72h. The competition between *E. coli* and *B. vulgatus* is exacerbated in certain CF-like conditions attributed to malabsorption, particularly high bile and high glycerol concentrations. To date, the role of glycerol in CF gut microbial interactions has not been explored. Through probing the role of glycerol in the competition model, we determined that the reduction in viability is dependent on live cells; cell-free supernatants of *E. coli* and *E. coli-B. vulgatus* co-cultures grown in glycerol do not recapitulate this reduction in viability. This finding may suggest the involvement of direct antagonism or acute mediators that act transiently.

We chose to take a genetic screening approach to identify *E. coli* genetic factors involved in this interaction. We generated a mariner transposon library in the CF-isolated *E. coli* strain 139H and performed a co-culture screen with a susceptible *B. vulgatus* strain (127O) in the presence of glycerol. Results from this screen definitively pointed to the biosynthesis of colibactin, a genotoxin associated with increased risk of colorectal cancer. *E. coli* mutants with transposon insertions in the *clb/pks* gene cluster rescue *B. vulgatus* growth in glycerol-supplemented co-cultures, independent of changes in *E. coli* growth. The involvement of colibactin production was further validated through the use of a *clbP* null mutant of *E. coli* as well as a chemical inhibitor of ClbP, both of which restored *B. vulgatus* viability in co-culture with glycerol.

Numerous mechanisms may explain the competition observed between *pks+ E. coli* and *B. vulgatus* strains. Firstly, there are a few lines of evidence that report variations in microbial composition attributed to colibactin production [65, 66]. Namely, mouse microbiomes with *pks+ E. coli* cluster separately from microbiomes lacking these microbes, and this separation is driven by a decrease in Firmicutes [66]. Functional analyses of the altered microbiomes indicate enhanced DNA repair pathways, suggesting the role of genotoxic activity on resident bacteria [66]. Additional studies support this finding: DNA damage is induced in distant microbes by *pks+ E. coli* in a contact-independent [67] and oxygen-dependent manner [65]. While our supernatant data do not agree with contact independence, it is likely that (1) temporal dynamics of colibactin’s stability and genotoxic activity are rather rapid and/or must be sustained for a duration longer than tested, and/or (2) the growth method in liquid media is not sufficient to recapitulate the contact independence, which was observed on soft agar.

In addition to the induction of DNA damage, there is evidence of the *clb/pks* island encoding not only colibactin, but also alternative intermediates and metabolites that are produced as derailment products of the NRPS-PKS [39, 41]. Prominently, the *pks* island is regulated by low iron levels; when iron is limited in the environment, expression of the biosynthetic cluster is induced via direct binding of ferric iron uptake regulator Fur to the promoter region of *clbA* [66]. When active, the pathogenicity island also influences the production of siderophores and microcin, which enhance the microbe’s iron-scavenging capabilities and subsequent colonization [41]. Siderophore production is critical to have a competitive advantage against pathogens, such as *Salmonella* Typhimurium [62, 68]. In addition, the *pks* pathway produces lipopeptides, including C12-Asn-GABA and C14-Asn [39]. C12-Asn-GABA is a known analgesic with pain-relieving activity, which may contribute to the probiotic activity of *pks+ E. coli* Nissle. C14-Asn has weak antimicrobial activity against *Bacillus subtilis* strains. Concurrently, through the extensive process of colibactin production, there is a large diversity of intermediate metabolites (e.g., precolibactins) that are produced, some of which at high abundances though with undetermined bioactivity [69]. It is likely that an abundance of alternative metabolites and intermediates are generated by this pathway, yet the environment that specifically dictates their regulation and the overall biological function of these alternative metabolites remain elusive.

Another mechanism by which *pks+ E. coli* may antagonize other microbes, including *B. vulgatus*, is through the induction of prophage. Literature suggests that prophage induction and subsequent lytic development are dependent on colibactin production and activity across genetically dissimilar bacteria [70]. Specifically, *pks+ E. coli* induces prophage in *Salmonella, Staphylococcus, Citrobacter, Enterococcus* and *pks-* strains of *E. coli*. Prophage induction not only activates lytic replication of the prophage and kills the bacterial cell, but also potentiates a phage epidemic within the broader microbial community [71]. Thus, this observation raises the possibility that *pks+* strains of *E. coli* kill *B. vulgatus* strains harboring resident prophage in our system, a finding that may have a wider array of ecological implications in vivo.

The role of glycerol in this interaction is unresolved yet points to the connection between the *pks* machinery and fatty acid metabolism [69]. It has been shown that some of the most abundant alternative metabolites produced by this machinery are gamma-lactam derivatives [69], suggesting crosstalk between colibactin and primary fatty acid biosynthesis. Considering this link, the enrichment of glycerol in the CF intestine may, in part, feed the production of *pks*-driven alternative metabolites. Further research is needed to determine the correlation between free glycerol levels and regulation of the *pks* pathway, colibactin and alternative metabolite production.

While the prodrug quantification results point to the trending increase in colibactin in CF stool, it is important to note that absolute concentration of colibactin in the context of the gut is unknown due to its extreme instability and recalcitrance to isolation [56]. Inference of colibactin activity via prodrug quantification and metagenomics analyses may shed light on its potential for production under certain physiologies, however these approaches do not confirm enzymatic activity and consequent genotoxicity. Colibactin’s instability in the presence of microaerophilic environments suggests its activity in the gut environment is restricted in space to anoxic regions and thus the toxin likely acts locally – this reasoning may explain the lack of toxicity by supernatants on *B. vulgatus* isolates. Therefore, bulk analysis of stool may miss the intricacies of colibactin activity in terms of in vivo microbial competition.

Here, we present findings that suggest a previously unexplored role of glycerol and potentially other breakdown products of fat catabolism in regulation of colibactin production. Because fat malabsorption is not exclusive to CF populations, the observation of colibactin-mediated microbial dysbiosis may be applicable across disease cohorts and highlight a potential focus for therapeutics.

## Materials and Methods

Detailed descriptions of strains, media, growth conditions, co-culture assays, linear regression, glycerol quantification, supernatant assays, mariner transposon mutagenesis, quantification of prodrug in bacterial cultures and stool, ClbP inhibitor assays, *E. coli* DNA extraction and whole genome sequencing, homology analysis, stool DNA isolation and sequencing, metagenomic sequencing analyses and all associated statistical analyses are provided in Supplementary Information Appendix (SI Appendix).

## Acknowledgments

We would like to thank the team at Michigan State University’s Mass Spectrometry and Metabolomics Core (MSU MSMC), including Cassandra Johnny, Anthony Schilmiller and James O’Keefe, for their contributions to this work. We also thank Dr. Christian Jobin (University of Florida, Gainesville, FL) for providing *E. coli* NC101 and *clbP* mutant strains. This study was funded by NIH P20GM130454 to Dartmouth’s Center for Quantitative Biology, NIH T32/HL134598 to KEB, CFF/OTOOLE23G0 and NIH/ES033988-01A1 to GAO.

