## Supplemental Information for "A genotoxin associated with colorectal cancer linked to gut dysbiosis in children with cystic fibrosis"

PNAS

*George A. O'Toole

**Author Contributions:** K.E.B. and G.A.O. designed research; K.E.B., S.V.S., A.V.B., C.E.M, M.A.A.R., R.A.V., and R.D.R. performed research; K.E.B., S.V.S., S.M.S., and M.A.A.R. analyzed data; S.M.S, M.A.A.R., and E.P.B. contributed new reagents/analytical tools; J.L.S., J.C.M., and G.A.O. obtained clinical samples; K.E.B. and G.A.O. wrote the paper.

**This PDF file includes:**

- Supplementary Materials and Methods
- Supplemental Figures 1 to 11
- Supplemental Tables 1, 3-10; Supplemental Tables 2, 11-12 attached separately
- Supplemental References

### Materials and Methods

#### Strains, media and growth conditions

Strains and plasmids used in this study are listed in **Supplemental Table 1**. *Bacteroides fragilis, thetaiotaomicron* and *vulgatus* were used throughout this study. *Bacteroides* spp. were routinely patched from frozen stocks on defibrinated blood sheep (Northeast Laboratories) agar plates supplemented with 100 μg/mL gentamicin then grown for 48h anaerobically at 37°C. A liquid culture of *Bacteroides* was grown in supplemented brain-heart infusion (BHIS) containing 37 g/L BHI (Becton Dickinson Ref.#237500), 1 g/L L-cysteine (VWR Cat.#0206), 10 mL/L hemin (100 mg/mL stock; Sigma-Aldrich Cat.#51280) and 0.2% final concentration of sodium bicarbonate (Fisher Scientific Cat.#S233) buffered to pH 7 using 100 mM MOPS (Sigma-Aldrich Cat.#M1254). *E. coli* strains were routinely patched from frozen stocks on Lysogeny broth (LB) agar plates (125 g/L yeast extract, 250 g/L tryptone, 125 g/L NaCl, 375 g/L agar), grown aerobically for 16h at 37°C, then a liquid culture was grown in LB liquid (100 g/L yeast extract, 200 g/L tryptone, 100 g/L NaCl) aerobically with agitation for 16h at 37°C. Medium was supplemented with 50 μg/mL carbenicillin and 62.5 μg/mL diaminopimelic acid (DAP) for the *E. coli* strain (SMC 9201) harboring the transposon-containing shuttle vector pBT20. Medium was supplemented with 15 μg/mL gentamicin for the *E. coli* Tn*M* mutants, or those with the transposon inserted into the genome.

Monoculture and co-culture assays performed between *B. vulgatus* and *E. coli* strains were grown in MiPro buffered to pH 7 with 100 mM MOPS [1] or MiPro supplemented with 8 g/L porcine stomach mucin (median-CF-MiPro concentration [2]), MiPro pH 7 + 1 g/L bile salts (low-CF-MiPro concentration), MiPro pH 7 + 2 g/L bile salts (median), MiPro pH 7 + 1% glycerol (median), MiPro pH 7 + 1 mM sodium nitrate (median), MiPro pH 7 + 1 mM sodium sulfate (median), MiPro pH 7 + 1 mM sodium formate (median), MiPro pH 7 + 1 μM hydrogen peroxide (low), MiPro pH 7 + 1 μg/mL and 10 μg/mL Bactrim (low and median, respectively), and MiPro pH 6 buffered with 100 mM MEPS. For all vendor and media preparation information, reference the methods in Barrack et al., 2024 [2]. Following monoculture and/or co-culture, *B. vulgatus* were selectively plated on blood sheep agar supplemented with 100 μg/mL gentamicin and 10 μg/mL nalidixic acid (counter-selection against *E. coli*) and grown anaerobically for 48h at 37°C; *E. coli* was selectively plated on LB agar plates and grown for 16h aerobically at 37°C.

#### Co-culture assay

*Bacteroides* spp. strains were grown on blood sheep agar plates supplemented with 100 μg/mL gentamicin for 48h prior to the experiment. *E. coli* strains were grown in 5 mL of LB at 37°C for ~16h with agitation. On the day of the experiment, each strain of *Bacteroides* and *E. coli* were normalized to an $OD_{600}$ = 0.05 and inoculated in a 1:1 ratio in each respective medium used: MiPro pH 7 and MiPro supplemented with each component, as listed in **Strains, media and growth conditions**. Each co-culture pair was coupled with a monoculture control of each strain used, also inoculated at a final $OD_{600}$ of 0.05. Co-cultures were incubated anaerobically for 72h at 37°C with use of the BD GasPak system (Cat. #260678). Following 72h incubation, growth of each strain was quantified by plating for CFU/mL. To do this, 72h cultures were 10-fold serially-diluted in sterile PBS then all dilutions were plated on selective medium for each species: blood sheep agar supplemented with 100 μg/mL gentamicin and grown anaerobically for 48h for *Bacteroides,* LB agar grown aerobically for 24h for *E. coli.*

Co-cultures of *B. vulgatus* and *E. coli* Tn*M* mutants follow a similar experimental design, with some alterations to selection. Specifically, *E. coli* Tn*M* mutants were grown in 5 mL LB supplemented with 15 μg/mL gentamicin for ~16h with agitation prior to experimental set-up. On the day of the experiment, methods are same as described above. Following 72h incubation, growth of each strain was quantified by plating for CFU/mL on selective medium: blood sheep agar supplemented with 100 μg/mL gentamicin (selection) and 10 μg/mL nalidixic acid (counter-selection) and grown anaerobically for 48h for *Bacteroides,* LB agar grown aerobically for 24h for *E. coli* WT and Tn*M.*

#### Linear regression

The $log_{10}$-transformed CFU/mL for *B. vulgatus* strains (n=6) and *E. coli* strains (n=8) grown in monoculture and co-culture in MiPro and individual CF-relevant components were compiled in **Supplemental Table 2**. Appropriate metadata including bacterial species, strain, competition status (monoculture vs. co-culture), the strain identification of the partner in co-culture, respective concentration of each tested CF-like feature (Base, Low, Median, No) and replicate were associated with the growth data. “Base” indicates levels used in MiPro, “Low” and “Median” indicate levels used in low- and median-CF-MiPro, respectively, and “No” indicates the complete absence of that feature. First, a linear model was performed to account for the contribution of species and competition status. Then, each species and competition status were subset to generate four separate metadata files: *B. vulgatus* monoculture, *B. vulgatus* co-culture, *E. coli* monoculture and *E. coli* co-culture. On each file, a linear model was performed to account for the contribution of each CF-like feature. Furthermore, strain-specific effects in low and median bile concentrations (Strain + Bile + Strain*Bile) were determined by performing a linear model on the *B. vulgatus* monoculture log-transformed CFU/mL. Output from the *B. vulgatus* mono- and co-culture linear models in the form of R-squared values and p-values were plotted as a bar plot using GraphPad Prism (version 10.4.2).

**Glycerol quantification**

Raw stool was weighed and thawed in individual 1.7 mL microtubes. Stool was homogenized using two 3mm ball bearings (VXB Cat.# Kit12064) and 1.2 mL of extraction solvent (2:1 chloroform:methanol (v/v), 0.01% (w/v) butylated hydroxytoluene (BHT, Sigma Cat.#218405) and 0.1% formic acid (Sigma, Cat.# FX0440)). The homogenate was vortexed then spun down at 13,000 *x g* for 10 min to precipitate any debris. The supernatant (800 μL) was transferred to a new microtube and vacuum concentrated at low speed using the Savant Speed Vac SC110 overnight. Samples were stored at -20°C until sent for further processing to Michigan State University Mass Spectrometry and Metabolomics Core (MSMC).

Once dried samples arrived at MSMC, samples were reconstituted in 800 μL of extraction solvent (2:1 chloroform:methanol, 0.1% formic acid), followed by addition of 500 μL of MilliQ water and mixing by vortex. Samples were centrifuged to separate phases and the upper aqueous phase was transferred to a new tube. Another 500 μL of MilliQ water was added to the organic phase, mixed and centrifuged and the aqueous phase was combined with the original aliquot. This combined aqueous phase was transferred to a new tube and used for free glycerol quantification.

A 50 μL aliquot of the combined aqueous phase was transferred to a new tube and 200 μL of acetonitrile and 5 uL of a 50 μg/ml glycerol-d5 internal standard solution was added. Samples were then analyzed using a Waters Acquity TQD tandem quadrupole mass spectrometer interfaced with a Waters Acquity UPLC. 10 μL of sample was injected onto a Waters Acquity UPLC BEH-Amide column (2.1x100 mm) and compounds were separated using the following gradient: initial conditions were 10% mobile phase A (1 mM ammonium acetate + 0.1% acetic acid in water) and 90% mobile phase B (acetonitrile) and held until 1.5 min, then ramp to 90% A at 3.5 min and hold until 4 min, return to 90 % B at 4.01 min and hold until 7 min. The column temperature was 35°C and flow rate was 0.3 mL/min. Compounds were ionized by electrospray ionization operating in negative ion mode. Parent ion and daughter ion masses were 151 and 59 for glycerol and 156 and 59 for glycerol-d5. A cone voltage of 10 and collision energy of 10 was used for both compounds. Concentrations were calculated using an external standard curve for glycerol using the Masslynx software.

Concentrations were then normalized to the raw weight of stool for each sample (μg/mg stool), plotted by genotype using GraphPad Prism (version 10.4.2). Associated metadata (e.g., genotype, age of the donor and batch of analysis) were assigned and a mixed effect linear model was performed (R package: lme4, version 1.1.34) [3]. The fixed variable in the linear model was genotype; the random variables were age of donor and batch.

**Supernatant assays**

*B. vulgatus* CFPLTA003-2B was grown on blood sheep agar plates supplemented with 100 μg/mL gentamicin for 48h prior to the experiment. *E. coli* 139H was grown in 5 mL of LB at 37°C for ~16h with agitation. On the day of the experiment, each strain of *Bacteroides* and *E. coli* were normalized to an $OD_{600}$ = 0.05 and inoculated in a 1:1 ratio in MiPro and MiPro+1% glycerol in a volume of 100 μL per well in flat-bottom 96-well plates (Corning Ref.#353072). The co-culture pair was coupled with a monoculture control of each strain, also inoculated at a final $OD_{600}$ of 0.05. Approximately 12 technical replicates were set up to reach a total volume of ~1.2 mL per biological replicate. Monocultures and co-cultures were grown anaerobically at 37°C for 72h with use of Coy Laboratory Products (2–3% $H_2$, 10% $CO_2$, $N_2$ balanced atmosphere). While maintaining an anoxic environment, the culture was centrifuged at 8,000 *x g* for 2 min and the supernatant was filter-sterilized using 0.22 μm filters. The resulting sterile supernatant was combined 1:1 with fresh de-gassed MiPro, then monocultures of *B. vulgatus* clinical isolates (n=5) and *E. coli* 139H were inoculated at a final $OD_{600}$ = 0.05. Growth controls include *B. vulgatus* or *E. coli* in 100% MiPro pH 7 or 100% MiPro pH 7 + 1% glycerol. After 24h of anaerobic growth, cell density was assessed via CFU/mL plating on sheep blood agar supplemented with 100 μg/mL gentamicin (for *B. vulgatus*) and LB agar (for *E. coli*). Statistical analysis was performed using 2-way ANOVA with Dunnett's multiple comparisons test comparing MiPro to all other conditions within each strain.

**Mariner transposon mutagenesis**

*E. coli* 139H WT and *E. coli* SMC 9201 (X7213, DAP-auxotroph, kanamycin-resistant, pir-positive) carrying the pBT20 plasmid were grown in LB and LB+62.5 μg/mL DAP+50 μg/mL carbenicillin overnight at 37°C, respectively, with agitation. Cell cultures were $OD_{600}$-normalized to 0.6, then cell pellets were centrifuged for 3 min at 3,610 x *g*, washed and resuspended in LB+62.5 μg/mL DAP. The mixture was plated in a 1:1 ratio on LB agar supplemented with 62.5 μg/mL DAP to facilitate conjugation of the pBT20 plasmid into *E. coli* 139H. Plates were incubated for 24h at 37°C, and the cell mixture was scraped up in 1 mL phosphate buffer saline (PBS), washed once with PBS, then dilutions were plated on LB agar supplemented with 15 μg/mL gentamicin to select for *E. coli* 139H mutants with chromosomal transposon insertions. DAP was not added to these selection plates as a form of counter-selection against the donor strain, SMC 9201. Individual candidates were picked with sterile pipette tips and inoculated in BHIS + 15 μg/mL gentamicin in sterile 96-well flat bottom plates, and the plates were incubated at 37°C for 16h. Each 96-well plate contained three control wells: (A1) *E. coli* 139H WT in monoculture, (A2) empty well (reserved for *B. vulgatus* 127O in monoculture control), (A3) *E. coli* 139H WT (and *B. vulgatus* 127O to be added later for co-culture control). After incubation, an equal volume, 100 μL, of 50% sterile glycerol was added to each well and plates were frozen at -80°C.

Mutants were then screened for their ability to reduce *B. vulgatus* 127O viability in glycerol-supplemented co-culture by using a 96-pin replicator (Dankar) to transfer inoculum from the frozen plates to a new sterile 96-well flat bottom plate containing BHIS, then grown overnight at 37°C. Concurrently, a culture of *B. vulgatus* 127O was grown on blood sheep agar supplemented with 100 μg/mL gentamicin and grown for 48h at 37°C anaerobically. The 127O culture was diluted to $OD_{600}$ = 0.05 and added to a fresh 96-well flat bottom plate containing sterile MiPro pH 7 + 1% glycerol. This normalized culture was added to wells A2-H12, reserving A1 for the *E. coli* 139H WT monoculture control. To this plate, the overnight cultures of the transposon mutants were replicated. The product of these transfers results in a plate containing MiPro pH 7 + 1% glycerol and a relatively equal inoculum of *E. coli* Tn*M* mutants and *B. vulgatus* 127O, with *E. coli* 139H WT monoculture, *B. vulgatus* 127O monoculture and respective co-culture control wells. This co-culture assay was incubated anaerobically for 72h, then each microbe's growth was quantified using selective media: blood sheep agar + 100 μg/mL gentamicin + 10 μg/mL nalidixic acid (anaerobic) to select for *B. vulgatus* and LB agar (aerobic) to select for *E. coli* WT and transposon mutants. The selective media plates were grown for 24h in the respective oxygen tensions, then growth was qualitatively addressed. Criteria to qualify as a candidate included: (1) viable *E. coli* growth on LB agar, (2) *B. vulgatus* growth visible minimally, partially or fully relative to Well A2, the *B. vulgatus* 127O monoculture control. Growth spots containing less than five countable colonies were considered to restore viability minimally; spots with five or more colonies were considered to restore viability partially; spots with full growth were considered to have full restoration, as shown in **Supplemental Figure 5**, bottom right. See **Supplemental Figure 5** for a schematic of the methods.

*E. coli* candidates that did not reduce *B. vulgatus* viability to any degree were picked into a sterile 96-well plate containing BHIS + 15 μg/mL gentamicin, grown overnight at 37°C, then glycerol-stocked frozen at -80°C to generate a sub-library of primary hits. The above screening process was repeated with primary hits to validate and determine secondary hits that showed no reduction in *B. vulgatus* 127O viability relative to monoculture. Validated candidates were then struck onto fresh LB agar with 15 μg/mL gentamicin, used to start an overnight LB+15μg/mL gentamicin liquid culture and glycerol-stocked at -80°C for future experimentation. From these frozen stocks, candidates were rescreened a third time in the typical co-culture assay with CFU/mL as the output to confirm full viability of *B. vulgatus*. The location of transposon insertion for secondary candidates was determined using arbitrary primed PCR as described by O'Toole et al. [4]. Based on these results, primers were designed to amplify and sequence the chromosomal flanks of the predicted insertion site to confirm the exact insertion location.

**Quantification of prodrug in bacterial cultures**

For 24h cultures:
All *E. coli* strains (Nissle, 143D, 139H, 139H *clbN::*Tn*M*, NC101 and NC101 Δ*clbP*) were grown in 5 mL of LB at 37°C for ~16h with agitation. On the day of the experiment, each strain of *E. coli* was normalized to an $OD_{600}$ = 0.05 and inoculated in MiPro and MiPro+1% glycerol in a volume of 100 μL per well in flat-bottom 96-well plates (Corning Ref.#353072). Approximately 12 technical replicates were set up to reach a total volume of ~1.2 mL per biological replicate. Cultures were grown anaerobically for 24h at 37°C with use of the Coy Laboratory Products (2–3% $H_2$, 10% $CO_2$, $N_2$ balanced atmosphere). Following 24h incubation, one technical replicate from each condition was plated for CFU/mL enumeration on LB agar, grown overnight aerobically at 37°C. The remaining 11 technical replicates were pooled in a 1.5 mL Eppendorf tube, then flash frozen at -80°C.

For 72h cultures:
*B. vulgatus* 127O was grown anaerobically on blood sheep agar plates supplemented with 100 μg/mL gentamicin for 48h prior to the experiment. *E. coli* 139H was grown in 5 mL of LB at 37°C for ~16h with agitation. On the day of the experiment, each strain was normalized to an $OD_{600}$ = 0.05 and inoculated in a 1:1 ratio in MiPro and MiPro+1% glycerol in a volume of 100 μL per well in flat-bottom 96-well plates (Corning Ref.#353072). The co-culture pair was coupled with a monoculture control of each strain, also inoculated at a final $OD_{600}$ of 0.05. Approximately 12 technical replicates were set up to reach a total volume of ~1.2 mL per biological replicate. Monocultures and co-cultures were grown anaerobically for 72h at 37°C with use of Coy Laboratory Products (2–3% $H_2$, 10% $CO_2$, $N_2$ balanced atmosphere). Following 24h incubation, one technical replicate from each condition was plated for CFU/mL enumeration on both LB agar (for *E. coli*) and blood sheep agar supplemented with 100 μg/mL gentamicin (for *B. vulgatus*). The remaining 11 technical replicates were pooled in a 1.5 mL Eppendorf tube, then flash frozen at -80°C.

500 µL of bacterial cultures were flash frozen and lyophilized until dryness. The samples were thawed on ice and 250 µL of extraction solution (MeOH:MeCN:$H_2O$, 2:2:1 ratio, containing 100 nM of D-27 myristoyl-prodrug internal standard) was added to each tube, and sonicated for five minutes. Samples were centrifuged (21,300 *g* x 15 min, 4 ºC), and the supernatant (200 µL) was carefully aspirated, and filtered through a 0.2 µm wwPTFE plate filter (Cytiva) into a 96-well plate. Samples were analyzed by UPLC–MS/MS on a Waters Xevo TQ-S UPLC-triple quadrupole with a Acquity UPLC H-Class System using a Cortecs UPLC C8 column (Waters corporation, 1.6µm, 2.1 mm x 75 mm). The conditions were as follows: 0.5 mL/min flow rate, column temperature: 40 ºC, 1 µL injection, 10% solvent B in solvent A for 0.5 min, a linear gradient increasing to 95% solvent B in solvent A over 0.5 min, holding 95% solvent B in solvent A for 1.0 min, followed by a linear gradient back to 10% solvent B in solvent A over 0.6 min, and re-equilibration at 10% solvent B in solvent A for 0.9 min (Solvent A, water + 0.1% formic acid; Solvent B, acetonitrile + 0.1% formic acid). The mass spectrometer was run in negative mode MRM, with the following conditions: capillary voltage, 2.1 kV, cone voltage, 4 V; source offset voltage, 50 V; desolvation temperature, 200 ºC; desolvation gas flow, 800 L/h; cone gas flow, 150 L/h; Nebulizer, 7.0 ba. Ions of interest were quantified by monitoring the transitions *m/z* 341.3 → *m/z* 226.3 (cone voltage 50 V, collision

energy 24 V, retention time 1.90 min) for *N*-myristoyl-D-Asn, and *m/z* 368.5 → *m/z* 253.3 (cone voltage 58 V, collision energy 28 V, retention time 1.91 min) for the deuterated *N*-myristoyl-D-Asn internal standard. Each sample was collected with two replicate injections, and with three independent triplicates. A calibration curve of an authentic standard of *N*-myristoyl-D-Asn (0 – 1000 nM, in duplicates) containing 100 nM of D-27 myristoyl-prodrug internal standard was used to find the concentration of *N*-myristoyl-D-Asn in samples. Data was analyzed using the TargetLynx software platform (Waters) and Microsoft Excel. Concentrations of prodrug were normalized to *E. coli* CFU/mL and plotted using GraphPad Prism (version 10.4.2). Statistical analysis was performed using 2-way ANOVA with Šídák's multiple comparisons test.

**Quantification of prodrug in stool samples**

Frozen stool samples were placed inside bashing-bead tubes (Bio-Rad), and kept on ice. 750 µL of extraction solution (MeOH:MeCN:$H_2O$, 2:2:1 ratio, containing 100 nM of D-27 myristoyl-prodrug internal standard) was added to each tube, bead-beated twice for 2 minutes at room temperature, with cooling on ice after each agitation (approx. 5 minutes). Samples were centrifuged (5,000 *g* x 5 min, 4 ºC), and supernatants transferred to a separate microcentrifuge tube and spun again (21,300 *g* x 15 min, 4 ºC). The supernatant (200 µL) was carefully aspirated and filtered through a 0.2 µm wwPTFE plate filter (Cytiva) into a 96-well plate. Samples were analyzed by UPLC–MS/MS on a Waters Xevo TQ-S UPLC-triple quadrupole with a Acquity UPLC H-Class System using a Cortecs UPLC C8 column (Waters corporation, 1.6µm, 2.1 mm x 75 mm). The conditions were as follows: 0.5 mL/min flow rate, column temperature: 40 ºC, 1 µL injection, 10% solvent B in solvent A for 0.5 min, a linear gradient increasing to 95% solvent B in solvent A over 0.5 min, holding 95% solvent B in solvent A for 1.0 min, followed by a linear gradient back to 10% solvent B in solvent A over 0.6 min, and re-equilibration at 10% solvent B in solvent A for 0.9 min (Solvent A, water + 0.1% formic acid; Solvent B, acetonitrile + 0.1% formic acid). The mass spectrometer was run in negative mode MRM, with the following conditions: capillary voltage, 2.1 kV, cone voltage, 4 V; source offset voltage, 50 V; desolvation temperature, 200 ºC; desolvation gas flow, 800 L/h; cone gas flow, 150 L/h; Nebulizer, 7.0 ba. Ions of interest were quantified by monitoring the transitions *m/z* 341.3 → *m/z* 226.3 (cone voltage 50 V, collision energy 24 V, retention time 1.90 min) for *N*-myristoyl-D-Asn, and *m/z* 368.5 → *m/z* 253.3 (cone voltage 58 V, collision energy 28 V, retention time 1.91 min) for the deuterated *N*-myristoyl-D-Asn internal standard. Each sample was collected with two replicate injections, and normalized to the premeasured weight of wet stool. A calibration curve of an authentic standard of *N*-myristoyl-D-Asn (0 – 1000 nM, in duplicates) containing 100 nM of D-27 myristoyl-prodrug internal standard was used to determine the concentration of *N*-myristoyl-D-Asn in samples. Data was analyzed using the TargetLynx software platform (Waters) and Microsoft Excel. Concentrations of prodrug were normalized to stool weight (mg) and plotted using GraphPad Prism (version 10.4.2). Statistical analysis was performed using a linear model (RStudio version 2024.09.1+394) to account for both genotype and batch effects on normalized prodrug concentrations.

**ClbP inhibitor assays**

The ClbP chemical inhibitor was obtained from the lab of Emily Balskus (Harvard University, Cambridge, MA) and was synthesized using a previously reported procedure [5]. The chemical name of the boronate precursor to the active inhibitor is: (*S*)-*N*-(3-amino-3-oxo-1-(4,4,5,5-tetramethyl-1,3,2-dioxaborolan-2-yl)propyl)-4-phenylbutanamide. The boronate precursor was initially suspended in 100% DMSO to a stock concentration of 10 mM and frozen at -20°C. On the day of each experiment, the stock solution was thawed and diluted to a 100 µM working stock in sterile water (1% DMSO). For each of the in vitro assays, a final concentration of 1 µM ClbP inhibitor was used in MiPro pH 7, with a 0.01% DMSO vehicle control. All co-culture assays in the presence of the inhibitor were performed the same as described above.

***E. coli* clinical isolate DNA extraction and whole genome sequencing**

Each *E. coli* clinical isolate was grown in 5 mL LB for 16h at 37°C with agitation. A volume of 500 µL of the grown culture was pelleted at 3,381 x *g* for 3 min. The supernatant was discarded and 300 µL of Cell Lysis Buffer (QIAGEN Mat. #1126462) was added to the pellet, then incubated at 80°C for 5 min. The pellet was put on ice for 1 min, then 1.5 µL of RNase A was added and incubated at 37°C for 30 min. The sample was then put on ice for 1 min and 100 μL of Protein Precipitation Buffer (QIAGEN Mat. #1045701) was added. The pellet was mixed well by vortexing. The sample was centrifuged for 3 min at max speed and the supernatant was transferred to a new tube, to which 300 μL cold isopropanol was added and mixed by pipetting. The DNA was centrifuged for 3 min at max speed, supernatant was discarded and the remaining pellet was washed twice with cold 70% ethanol. The pellet was air-dried for ~10 min, then 50 μL molecular-grade water was added. To increase DNA yield, the resuspended DNA was incubated at 65°C for 1h, then DNA concentration and quality was measured using a Nanodrop.

Extracted genomic DNA (gDNA) was sequenced using Illumina NextSeq2000 (SeqCoast Genomics, Portsmouth, NH) producing 2x150 bp paired-reads. The *E. coli* short-read genomes were assessed for quality using FastQC (version 0.12.1), then the adapter sequences and poor quality reads were removed using Trimmomatic (version 0.39). The filtered reads were assembled de novo using Spades (version 3.15.5). The assembled genome was annotated using Prokka (version 1.14.6). The quality of the genome assemblies was evaluated using Quast (version 5.2.0) and BUSCO (version 5.3.2.). The *E. coli* genomes are available in BioProject PRJNA833080.

#### *E. coli pks/clb* homology analysis

Assembled *E. coli* genome protein sequences were first used to build a BLAST+ database for each genome using the makeblastdb command. The *E. coli* Nissle 1917 genome file was accessed via NCBI (Reference sequence: NZ_CP082949.1). In this strain, the annotated Clb sequences, of which all were detected, were used as the reference (query) sequence for multiple sequence alignment of each respective *E. coli* clinical isolate database using the blastp command. Hits were then filtered by e-value and percent identity, where a valid hit maintained both an e-value of less than or equal to 1E-10 and a percent identity to the query sequence of greater than or equal to 50%. Results are reported in **Supplemental Table 12**, containing the following parameters: query accession (*clb* gene in Nissle), subject sequence ID, expect value (e-value), Bit score, percent identity, query coverage per subject, and start/end positions of alignment in query and subject sequences. The top percent identity values of each subject sequence compared to Nissle (query) were plotted as a heatmap using GraphPad Prism (version 10.4.2). Percent identify values of 0% indicate sequences that did not meet the filtering requirements and were thus determined absent from the respective *E. coli* strain.

#### Metagenomic sequencing analyses: taxonomic profiling and functional pathway analysis

Metagenomic sequences are publicly available through BioProjects PRJNA1244851 [6] and PRJNA955235 [7]. Reads were filtered and trimmed when applicable using KneadData (version 0.12.0), trimming bases using default settings and Phred-33 quality scoring and removing reads that mapped to the human genome (GRCh38). After trimming, the average number of reads was 30,891,665 and 100% of samples had at least 23 million reads. Taxonomic assignment was performed using MetaPhlAn (version 4.0.6) with the mpa_vOct22_CHOCOPhlAnSGB_202212 database. Gene family and pathway assignment was performed using HUMAnN (version 3.7) with the full chocophlan.v201901_v31 and uniref90_annotated_v201901b_full databases. Reads were mapped to the UniRef_90 database, which represents gene families clustered at 90% identity. Clustering at 90% identity enables assignment of taxonomic groups for most gene families. Counts for each gene family are first normalized for gene length with reads per kilobase and then normalized for library depth with total sum scaling to enable comparison between samples. Lastly, differential abundance testing across conditions of interest was performed with MaAslin (version 1.14.1). Statistical significance of differentially abundant taxa and gene families was addressed through linear modeling (R package: lme4 version 1.1.34).

### Supplemental Figures

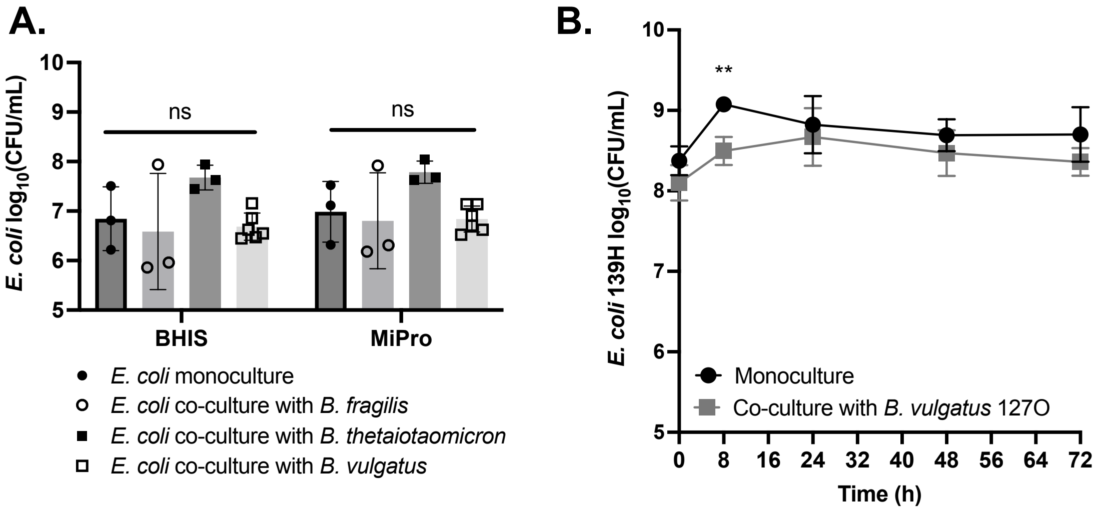

**Supplemental Figure 1. *E. coli* growth does not change between monoculture and co-culture with *Bacteroides* spp. (A)** *E. coli* 139H log-transformed CFU/mL in monoculture (filled circles) and co-culture with *B. fragilis* (open circles), *B. thetaiotaomicron* (filled squares) and *B. vulgatus* (open squares) in BHIS pH 7 and MiPro pH 7. Statistical analysis performed using 2-way ANOVA with Dunnett's multiple comparisons across growth status within each medium. **(B)** *E. coli* 139H log-transformed CFU/mL in monoculture (black) or co-culture with *B. vulgatus* 127O (grey) over the course of 72h in MiPro pH 7. Statistical analysis performed using 2-way ANOVA with Šídák's multiple comparisons at each time point (**, $p < 0.01$).

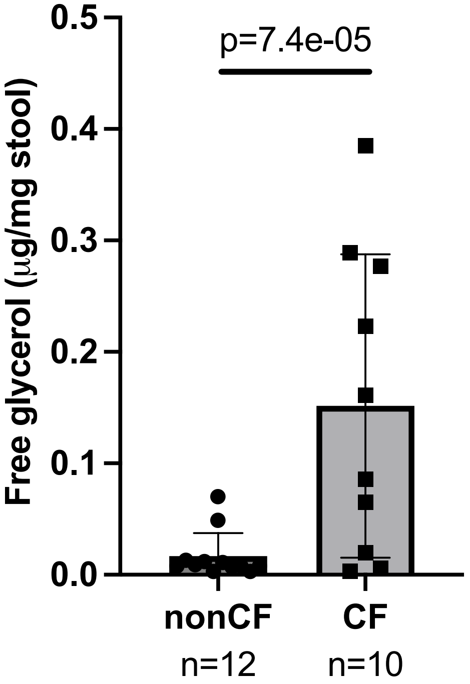

**Supplemental Figure 2. Free glycerol is detected at higher concentrations in CF stool compared to nonCF stool.** Free glycerol, or glycerol that is un-esterified to triacylglycerol, was quantified from fecal extracts of nonCF (n=12) and CF (n=10) stool from cwCF. Concentrations were normalized to weight of stool (mg). Statistical analysis performed using mixed effect linear modeling with age of donor and batch as the random variables, and genotype as the fixed variable.

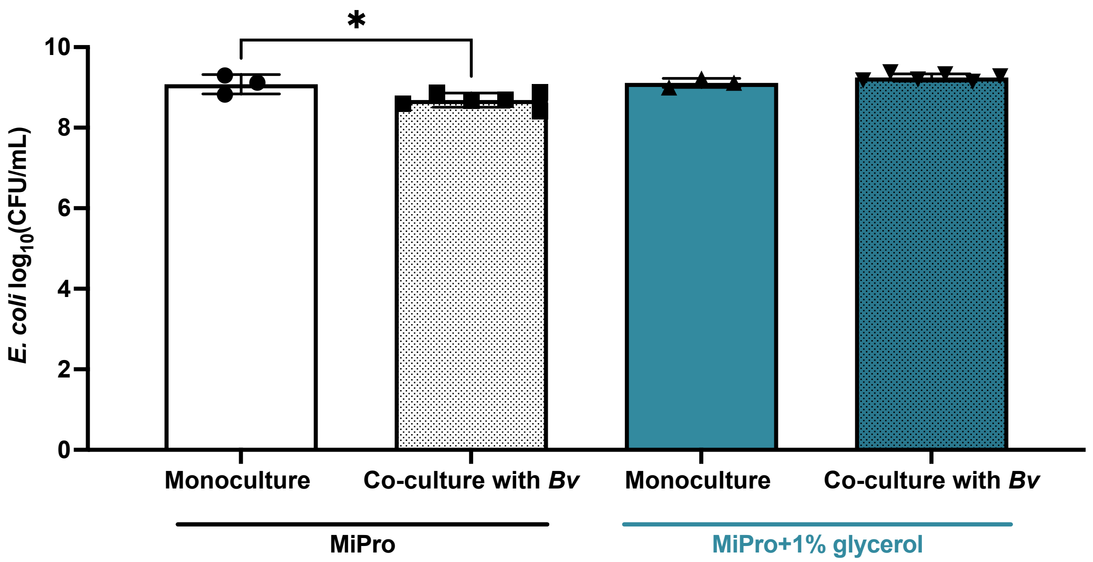

**Supplemental Figure 3. *B. vulgatus* viability reduction by *E. coli* 139H is independent of enhanced *E. coli* growth.** *E. coli* 139H log-transformed CFU/mL in monoculture (solid bars) and co-culture with *B. vulgatus* strains (n=6; patterned bars) in MiPro pH 7 (grey) and MiPro pH 7 + 1% glycerol (blue) for 72h. Statistical analysis performed using one-way ANOVA with Tukey's multiple comparisons; only significant ($P<0.05$) comparisons shown.

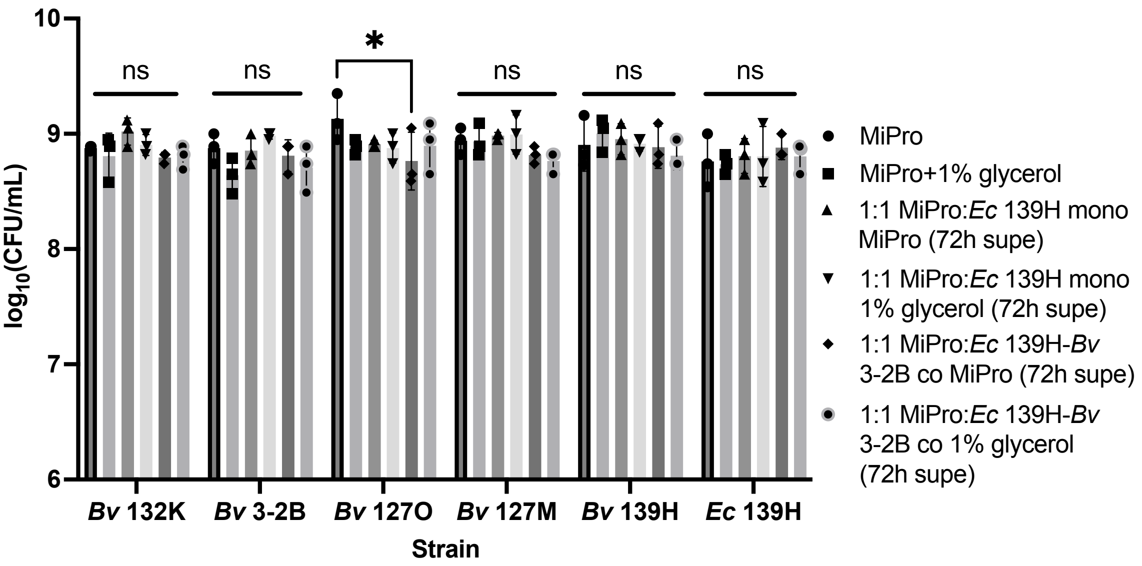

**Supplemental Figure 4.** Anoxic monoculture supernatants from (1) *E. coli* 139H grown in MiPro pH 7 (upright triangle) and MiPro pH 7+1% glycerol (inverted triangle) for 72h and (2) *E. coli* 139H and *B. vulgatus* CFPLTA003-2B grown in MiPro (diamond) or MiPro+1% glycerol (concentric circle) for 72h were collected and filter-sterilized. These anoxic cell-free supernatants were diluted in fresh MiPro and exposed to *B. vulgatus* strains (n=5) and *E. coli* 139H in monoculture. Controls include *B. vulgatus* strains (n=5) and *E. coli* 139H grown in monoculture in MiPro pH 7 (circle) or MiPro+1% glycerol alone (square) for 24h at 37°C under anaerobic conditions. Growth was quantified via CFU/mL enumeration on sheep blood agar supplemented with 100 μg/mg gentamicin for *B. vulgatus* and LB agar for *E. coli*. Statistical analysis performed using 2-way ANOVA with Dunnett's multiple comparisons across growth conditions within each bacterial strain (*, $p < 0.05$).

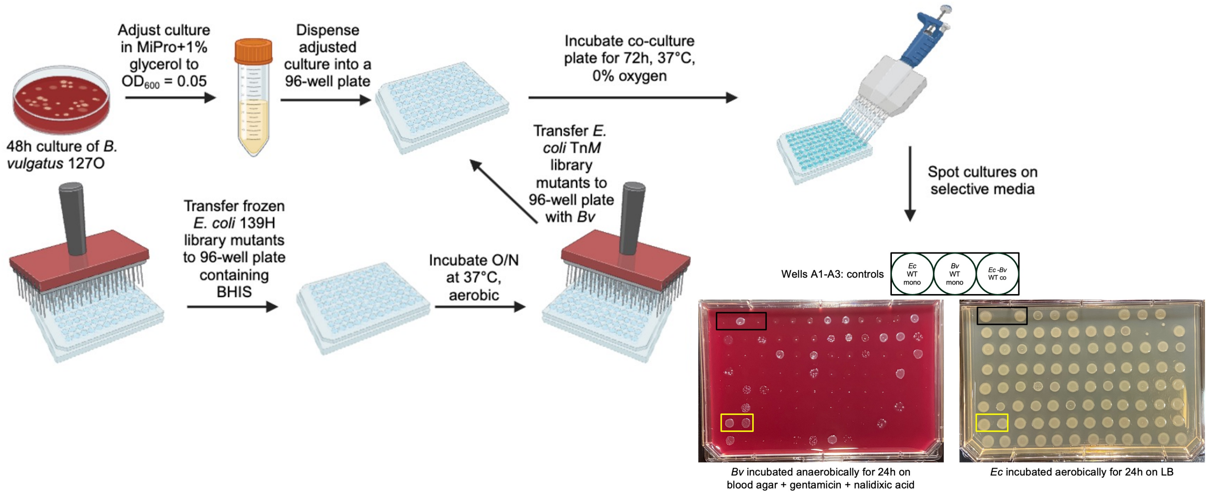

**Supplemental Figure 5. Schematic of mariner transposon screen.** On Day -2, *B. vulgatus* 127O was struck on blood sheep agar supplemented with 100 μg/mL gentamicin from a frozen stock and incubated at 37°C, anaerobically for 48h. On Day -1, frozen libraries of *E. coli* 139H Tn*M* were replicated into a sterile 96-well plate containing BHIS and incubated at 37°C, aerobically for 16h. On Day 0, the *B. vulgatus* 127O plates were scraped using a sterile pipette tip and $OD_{600}$-normalized to 0.05 in MiPro pH 7 supplemented with 1% glycerol. A volume of 100 μL was dispensed into each well (Wells A2-H12) of a sterile 96-well plate. Well A1 received only MiPro+1% glycerol. A volume of 3 μL of the overnight culture of the replicated *E. coli* Tn*M* library was transferred into wells A1, A3-H12 of the aforementioned plate. The co-culture was incubated at 37°C anaerobically for 72h. Following the incubation, the wells were mixed by pipetting and 3 μL was directly plated on two selective media: blood sheep agar supplemented with 100 μg/mL gentamicin and 10 μg/mL nalidixic acid, anaerobic growth for 24h for *B. vulgatus* 127O (“*Bv* 127O”) and LB agar, aerobic growth for 24h for *E. coli* WT and Tn*M.* Following 24h incubation, growth in control wells (A1-A3) were assessed for contamination. Proper control growth is shown above, whereby growth of *B. vulgatus* 127O is visible in its monoculture condition (A2), but not in the co-culture with *E. coli* 139H WT (A3), as reported in Figure 2. In addition, *E. coli* 139H WT growth is observed in its monoculture (A1) and co-culture conditions (A3). Spots of growth in wells A4-H12 were compared to the respective monoculture controls (A1-A3) to assign respective *B. vulgatus* restoration status. The highlighted boxes in yellow denote wells containing viable *E. coli* Tn*M* mutants that restored *B. vulgatus* 127O to its full extent, relative to its monoculture control (well A2). Figure made in BioRender.

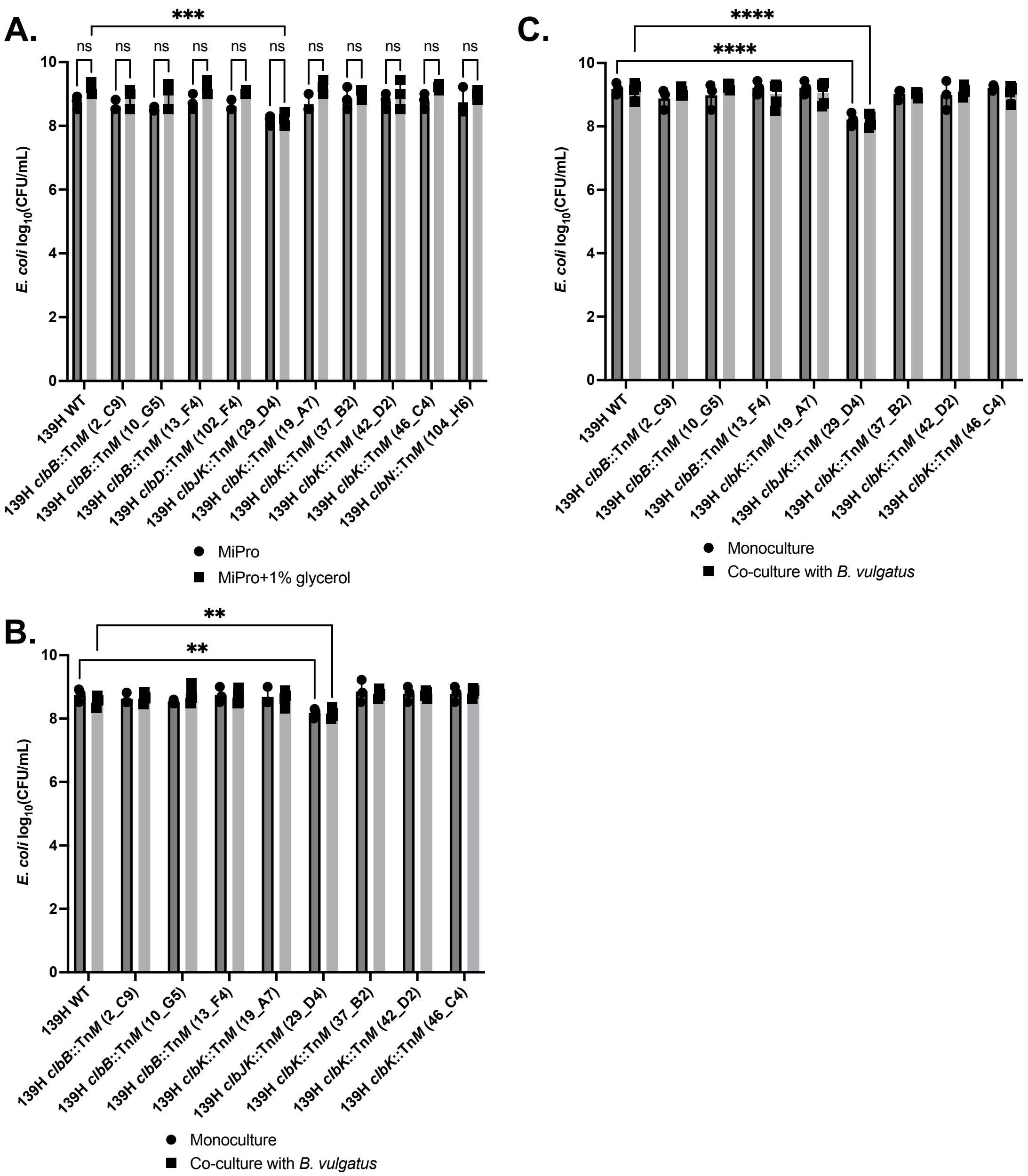

**Supplemental Figure 6. Restoration of *B. vulgatus* viability by *E. coli* Tn*M* strains is largely independent of differential *E. coli* growth. (A)** Monoculture growth of *E. coli* 139H WT and representative Tn*M* mutants in MiPro pH 7 and MiPro pH 7 + 1% glycerol for 72h anaerobically. Statistical analysis performed using 2-way ANOVA with Šídák's multiple comparisons (***, $p < 0.005$). **(B-C)** Endpoint (72h) log-transformed CFU/mL of *E. coli* 139H WT and representative *clb*::Tn*M* mutants in monoculture (circles) and co-culture with *B. vulgatus* (n=5; squares) in **(B)** MiPro pH 7 and **(C)** MiPro pH 7 + 1% glycerol. Statistical analysis performed using 2-way ANOVA with Šídák's multiple comparisons between each transposon mutant and the WT control for each growth condition (*, $p < 0.05$; **, $p < 0.01$; ***, $p < 0.005$; ****, $p < 0.001$).

A.

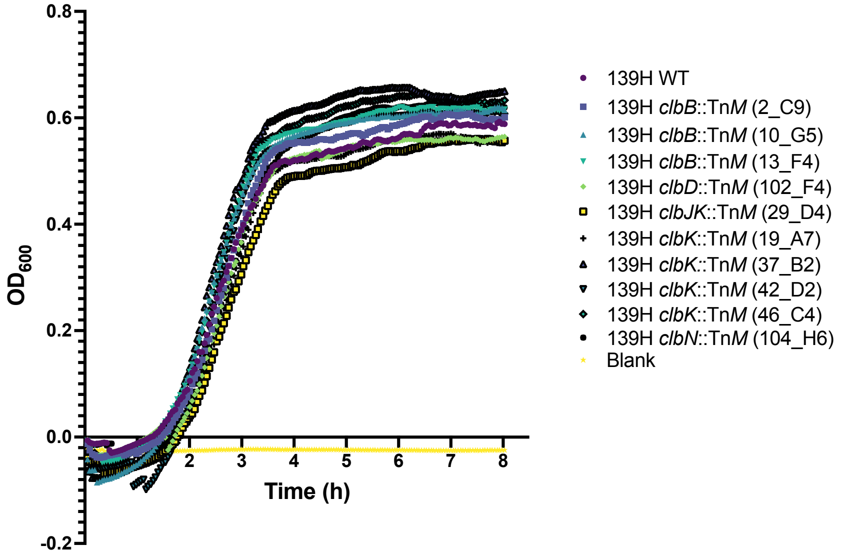

B.

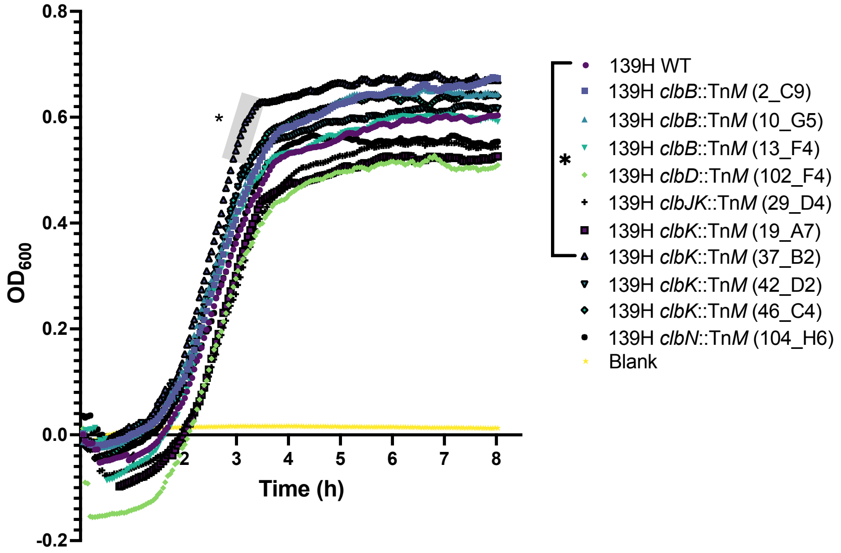

**Supplemental Figure 7. Growth kinetics of *clb*::Tn*M* mutants compared to *E. coli* 139H WT.** *E. coli* 139H WT and the representative *clb*::Tn*M* mutants were inoculated in **(A)** BHIS pH 7 and **(B)** BHIS pH 7 + 1% glycerol at an $OD_{600}$ = 0.01, then grown for 8h anaerobically at 37°C. Growth, as quantified by $OD_{600}$, is plotted against time (hours). Statistical analysis performed using 2-way ANOVA with Dunnett's multiple comparisons between each transposon mutant and the WT strain at each timepoint (*, $p < 0.05$). Error bars not shown for clarity.

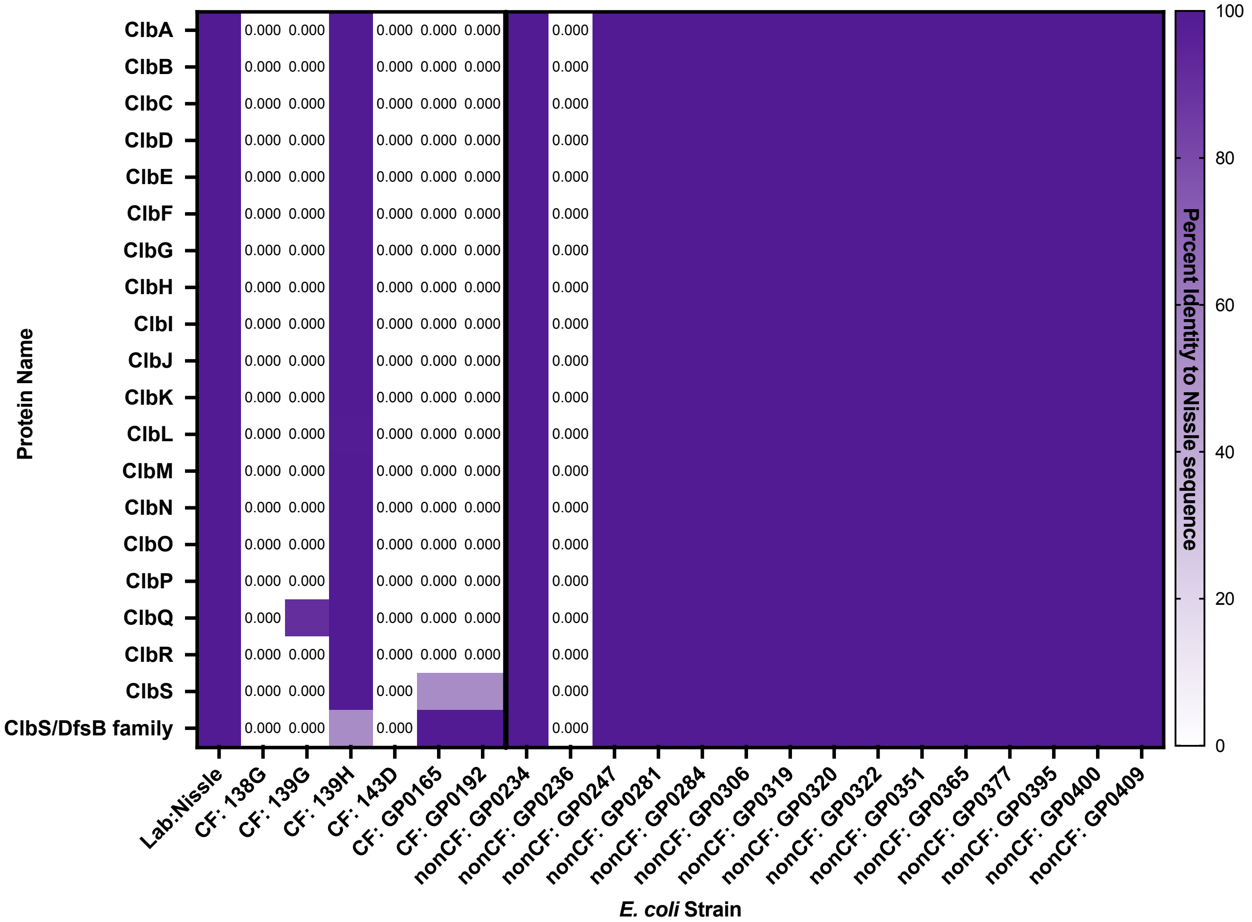

**Supplemental Figure 8. Sequence homology of the *clb* island across *E. coli* clinical isolates.** Protein sequence identity of all Clb proteins (Y-axis) in CF and nonCF clinical isolates of *E. coli* (X-axis), with *E. coli* Nissle 1917 protein sequences as reference. Hits were filtered by e-value and percent identity, where a valid hit maintained both an e-value of less than or equal to 1E-10 and a percent identity to the query sequence of greater than or equal to 50%. The top percent identity values of each subject sequence compared to Nissle (query) were plotted as a heatmap. Percent identify values of 0% indicate sequences that did not meet the filtering requirements and were thus determined absent from the respective *E. coli* strain.

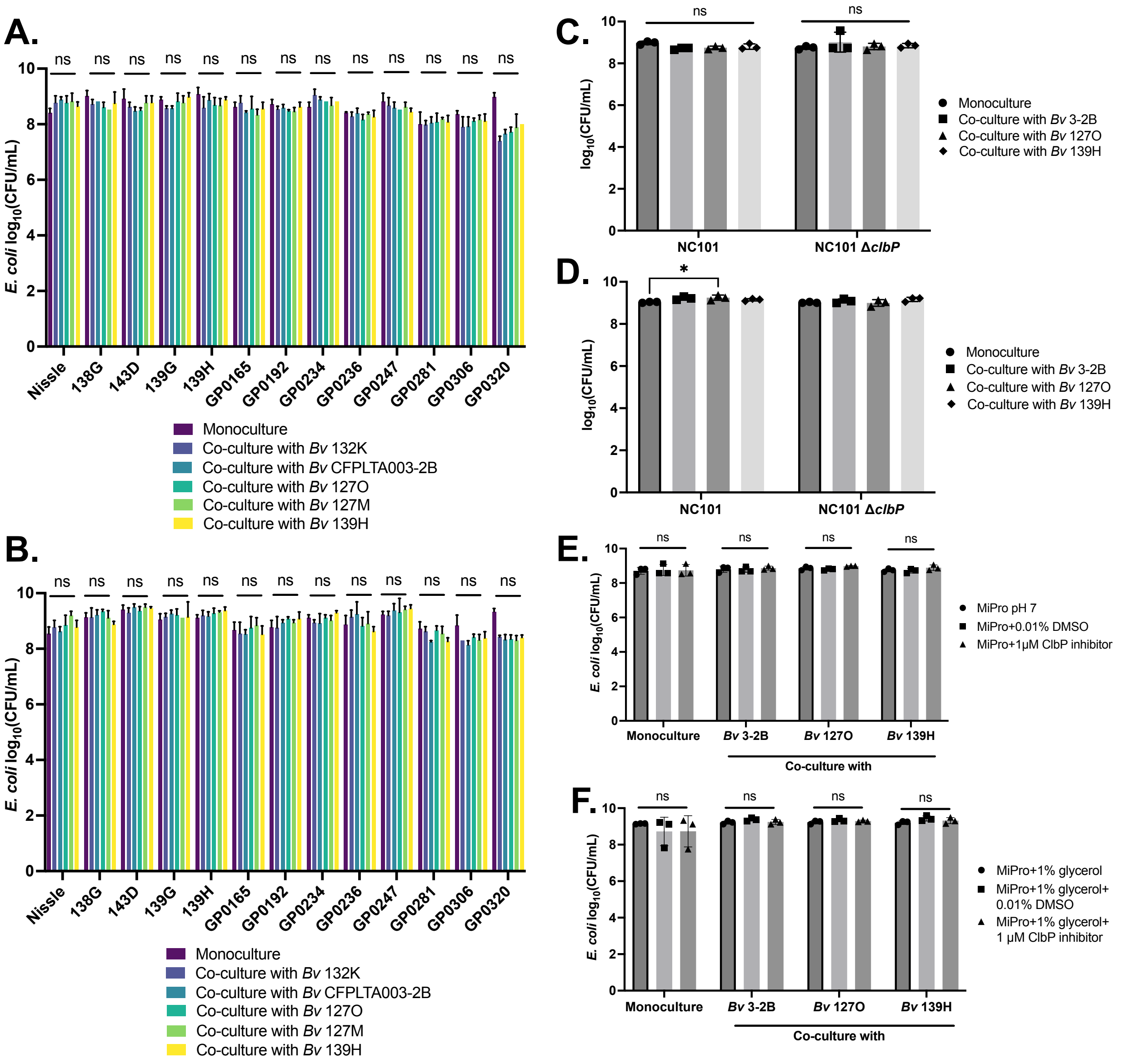

**Supplemental Figure 9. *E. coli* growth in monoculture and co-culture with *B. vulgatus* in MiPro pH 7 and MiPro pH 7 supplemented with 1% glycerol. (A-B)** *E. coli* strains (n=13) grown in monoculture (purple bars) and co-culture with 5 individual *B. vulgatus* isolates in **(A)** MiPro pH 7 and **(B)** MiPro pH 7 + 1% glycerol for 72h. Statistical analysis performed using 2-way ANOVA with Tukey's multiple comparisons between each co-culture condition to its respective monoculture control (ns, $p > 0.05$). **(C-D)** *E. coli* NC101 and NC101 ∆*clbP* grown in monoculture (circle) and co-culture with 3 individual *B. vulgatus* isolates: 3-2B (square), 127O (triangle), 139H (diamond) in **(C)** MiPro pH 7 and **(D)** MiPro pH 7 supplemented with 1% glycerol. Statistical analysis performed using 2-way ANOVA with Tukey's multiple comparisons between all growth conditions within each *E. coli* strain (*, $p<0.05$). **(E-F)** *E. coli* 139H grown in monoculture (circle) and co-culture with 3 individual *B. vulgatus* isolates: 3-2B, 127O, 139H in **(E)** MiPro pH 7 (circle), MiPro+0.01% DMSO (square), MiPro+1µM ClbP inhibitor (triangle) and **(F)** MiPro pH 7+1% glycerol (circle), MiPro pH 7+1% glycerol+0.01% DMSO (square), MiPro pH 7+1% glycerol+1µM ClbP inhibitor (triangle). Statistical analysis performed using 2-way ANOVA with Tukey's multiple comparisons between all growth conditions.

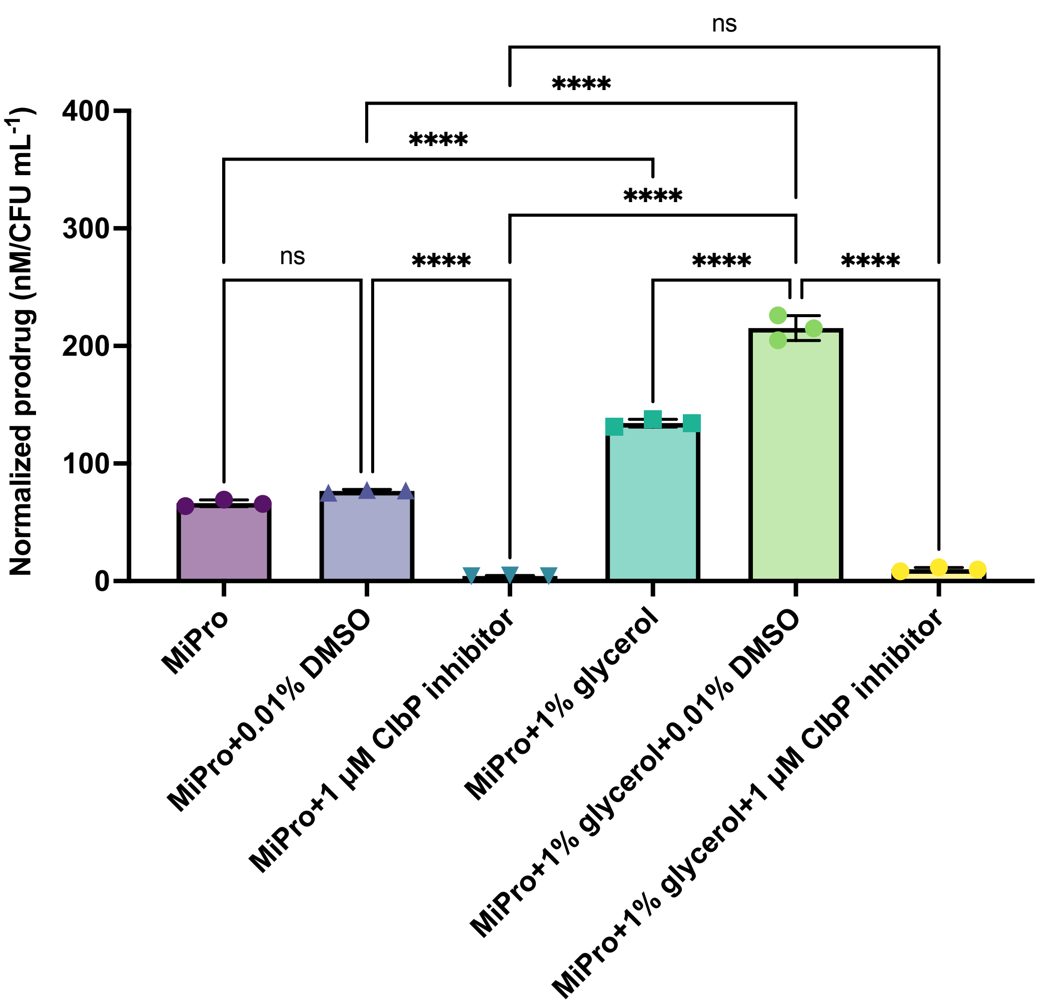

**Supplemental Figure 10. Colibactin-derived prodrug (*N*-myristoyl-D-Asn) production is reduced following treatment with a small molecule inhibitor of ClbP.** *E. coli* 139H was grown in monoculture anaerobically at 37°C for 72h in MiPro pH 7 (purple circle), MiPro pH 7+0.01% DMSO (upright triangle), MiPro pH 7+ 1 µM ClbP inhibitor (inverted triangle), MiPro pH 7+ 1% glycerol (square), MiPro pH 7+ 1% glycerol+0.01% DMSO (green circle) and MiPro pH 7+ 1% glycerol+ 1 µM ClbP inhibitor (yellow circle). Whole cultures were both plated for CFU/mL enumeration on LB agar and used for prodrug quantification via UPLC–MS/MS. Prodrug concentrations were normalized to cell count (nM/CFU $mL^{-1}$). Statistical analysis performed using one-way ANOVA with Šídák’s multiple comparisons (****, $p<0.001$).

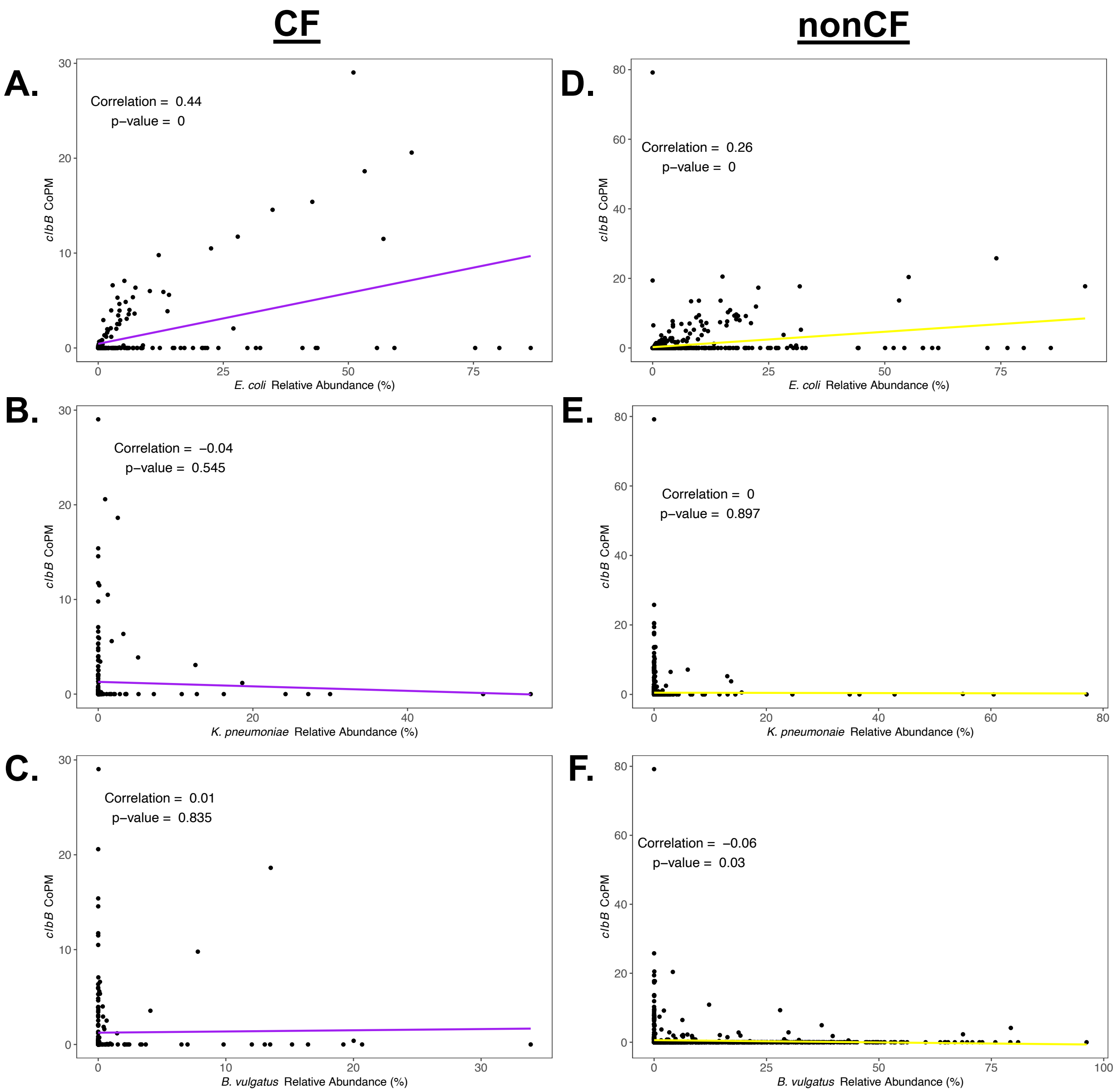

**Supplemental Figure 11. *clbB* gene count and species correlation plots in CF and nonCF stool metagenomes.** The abundance of *clbB* gene counts (copies per million reads: CoPM) are plotted against **(A, D)** *E. coli*, **(B, E)** *K. pneumonaie*, **(C,F)** *B. vulgatus* relative abundance (%) for **(A-C)** CF stool metagenomes (purple trend line, n=208) and **(D-F)** nonCF stool metagenomes (yellow trend line, n=2,560). Statistical analysis was performed using Pearson correlation.

### Supplemental Tables

**Supplemental Table 1. Bacterial strains and plasmids used in this study**

| Strain/plasmid | Genotype/Description | Reference/Source |
|---|---|---|
| ***Bacteroides fragilis*** | | |
| 126T-3B | isolated from stool from a CF child | [8] |
| 139C | isolated from stool from a CF child | [8] |
| CFPLTA004-1B | isolated from stool from a nonCF child | [8] |
| ***Bacteroides thetaiotaomicron*** | | |
| VPI-5482 | lab strain | [9] |
| 135X-1B | isolated from stool from a CF child | this study |
| GP1029 | isolated from a colonoscopy sample from a nonCF adult | this study |
| ***Phocaeicola vulgatus* (formerly *Bacteroides vulgatus*)** | | |
| 132K-1B | isolated from stool from a CF child | [8] |
| CFPLTA002-1B | isolated from stool from a nonCF child | [8] |
| CFPLTA003-2B | isolated from stool from a nonCF child | [8] |
| 127O | isolated from stool from a CF child | [8] |
| 127M | isolated from stool from a CF child | this study |
| b139H | isolated from stool from a CF child | [8] |
| ***Escherichia coli*** | | |
| Nissle 1917 | probiotic strain | [10] |
| 138G | isolated from stool from a CF child | this study |
| 143D | isolated from stool from a CF child | this study |

| | | |
|---|---|---|
| 139G | isolated from stool from a CF child | this study |
| e139H | isolated from stool from a CF child | this study |
| GP0165 | isolated from a colonoscopy sample from a CF adult (ID:GIECF100) | this study |
| GP0192 | isolated from a colonoscopy sample from a CF adult (ID:GIECF100) | this study |
| GP0234 | isolated from a colonoscopy sample from a nonCF adult (ID:GIEHV103) | this study |
| GP0236 | isolated from a colonoscopy sample from a nonCF adult (ID:GIEHV103) | this study |
| GP0247 | isolated from a colonoscopy sample from a nonCF adult (ID:GIEHV103) | this study |
| GP0281 | isolated from a colonoscopy sample from a nonCF adult (ID:GIEHV103) | this study |
| GP0284 | isolated from a colonoscopy sample from a nonCF adult (ID:GIEHV103) | this study |
| GP0306 | isolated from a colonoscopy sample from a nonCF adult (ID:GIEHV103) | this study |
| GP0319 | isolated from a colonoscopy sample from a nonCF adult (ID:GIEHV103) | this study |
| GP0320 | isolated from a colonoscopy sample from a nonCF adult (ID:GIEHV103) | this study |
| GP0322 | isolated from a colonoscopy sample from a nonCF adult (ID:GIEHV103) | this study |

| | | |
|---|---|---|
| GP0351 | isolated from a colonoscopy sample from a nonCF adult (ID:GIEHV103) | this study |
| GP0365 | isolated from a colonoscopy sample from a nonCF adult (ID:GIEHV103) | this study |
| GP0377 | isolated from a colonoscopy sample from a nonCF adult (ID:GIEHV103) | this study |
| GP0395 | isolated from a colonoscopy sample from a nonCF adult (ID:GIEHV103) | this study |
| GP0400 | isolated from a colonoscopy sample from a nonCF adult (ID:GIEHV103) | this study |
| GP0409 | isolated from a colonoscopy sample from a nonCF adult (ID:GIEHV103) | this study |
| SMC 9201 | X7213, DAP auxotroph, kanamycin-resistant, pir positive | [11] |
| NC101 | isolated from a mouse; carcinogeneic | [12] |
| NC101 Δ*clbP* | mutagenized using lamba-red recombinase to generate a null mutant of *clbP* in the parent NC101 background | [12] |
| **Plasmid** | | |
| SMC 1210 | pBT20 | [13] |

**Supplemental Table 2. Summarized growth data.** Attached separately due to size.

**Supplemental Table 3. Contribution of species and microbial competition on all CFU/mL.**

| **Predictors** | **Estimates** | **Confidence interval** | **p-value** |
|---|---|---|---|
| (Intercept) | 2.64 | 2.40-2.88 | **<0.001** |
| Species | 0.9 | 0.70-1.09 | **<0.001** |
| Co-culture | -0.35 | (-)0.6 - (-)0.1 | **0.007** |
| Observations | 2375 | | |
| R-squared/R-squared adjusted | 0.036 / 0.035 | | |

**Supplemental Table 4. *E. coli* CFU/mL in monoculture across CF-like conditions.**

| **Predictors** | **Estimates** | **Confidence interval** | **p-value** |
|---|---|---|---|
| (Intercept) | 3.14 | 3.08-3.2 | **<0.001** |
| Mucin [Median] | 0.13 | 0.04-0.22 | **0.003** |
| Bile [Median] | 0.16 | 0.07-0.25 | **0.001** |
| Glycerol [Median] | 0.14 | 0.05-0.23 | **0.002** |
| Nitrate [Median] | 0.14 | 0.05-0.23 | **0.003** |
| Sulfate [Median] | 0.13 | 0.03-0.22 | **0.007** |
| Formate [Median] | 0.12 | 0.02-0.21 | **0.014** |
| H2O2 [Low] | -0.18 | (-)0.37-0.01 | 0.07 |
| pH [Median] | 0.09 | (-)0.28-0.11 | 0.378 |
| Antibiotics [Low] | -0.15 | (-)0.34-0.05 | 0.135 |
| Antibiotics [Median] | -0.17 | (-)0.36-0.02 | 0.088 |
| Observations | 172 | | |
| R-squared/R-squared adjusted | 0.226 / 0.178 | | |

**Supplemental Table 5.** ***E. coli*** **CFU/mL in co-culture with** ***B. vulgatus*** **across CF-like conditions.**

| **Predictors** | **Estimates** | **Confidence interval** | **p-value** |
|---|---|---|---|
| (Intercept) | 3.25 | 3.23-3.27 | **<0.001** |
| Mucin [Median] | 0.03 | (-)0.01-0.06 | 0.103 |
| Bile [Median] | 0.03 | 0-0.07 | **0.049** |
| Glycerol [Median] | 0.06 | 0.02-0.09 | **0.001** |
| Nitrate [Median] | -0.02 | (-)0.06-0.01 | 0.177 |
| Sulfate [Median] | -0.01 | (-)0.05-0.02 | 0.499 |
| Formate [Median] | -0.01 | (-)0.04-0.03 | 0.67 |
| H2O2 [Low] | -0.26 | (-)0.33-(-)0.2 | **<0.001** |
| pH [Median] | -0.3 | (-)0.37-(-)0.23 | **<0.001** |
| Antibiotics [Low] | -0.33 | (-)0.4-(-)0.26 | **<0.001** |
| Antibiotics [Median] | -0.31 | (-)0.38-(-)0.24 | **<0.001** |
| Observations | 979 | | |
| R-squared/R-squared adjusted | 0.273 / 0.266 | | |

**Supplemental Table 6.** ***B. vulgatus*** **CFU/mL in monoculture across CF-like conditions.**

| **Predictors** | **Estimates** | **Confidence interval** | **p-value** |
|---|---|---|---|
| (Intercept) | 3.11 | 2.12-4.11 | **<0.001** |
| Mucin [Median] | 0 | (-)1.29-1.29 | 0.999 |
| Bile [Low] | -0.11 | (-)1.63-1.42 | 0.891 |
| Bile [Median] | -5.17 | (-)6.55-(-)3.78 | **<0.001** |
| Glycerol [Median] | -0.03 | (-)1.46-1.39 | 0.965 |
| Nitrate [Median] | 0.08 | (-)1.47-1.63 | 0.919 |
| Sulfate [Median] | 0.06 | (-)1.49-1.61 | 0.942 |
| Formate [Median] | 0.05 | (-)1.5-1.6 | 0.946 |
| H2O2 [Low] | -0.23 | (-)1.75-1.3 | 0.768 |
| pH [Median] | -0.14 | (-)1.67-1.38 | 0.855 |
| Antibiotics [Low] | -0.21 | (-)1.73-1.32 | 0.79 |
| Antibiotics [Median] | -0.12 | (-)1.65-1.4 | 0.873 |
| Observations | 250 | | |
| R-squared/R-squared adjusted | 0.294 / 0.262 | | |

**Supplemental Table 7.** ***B. vulgatus*** **CFU/mL in co-culture with** ***E. coli*** **across CF-like conditions.**

| **Predictors** | **Estimates** | **Confidence interval** | **p-value** |
|---|---|---|---|
| (Intercept) | 3.09 | 2.54-3.65 | **<0.001** |
| Mucin [Median] | 0.04 | (-)0.77-0.86 | 0.919 |
| Bile [Median] | -3.36 | (-)4.17-(-)2.55 | **<0.001** |
| Glycerol [Median] | -3.15 | (-)3.93-(-)2.37 | **<0.001** |
| Nitrate [Median] | -0.03 | (-)0.84-0.79 | 0.947 |
| Sulfate [Median] | -0.02 | (-)0.84-0.8 | 0.962 |
| Formate [Median] | -0.02 | (-)0.83-0.8 | 0.97 |
| H2O2 [Low] | -0.33 | (-)2.01-1.35 | 0.704 |
| pH [Median] | -0.29 | (-)1.97-1.39 | 0.736 |
| Antibiotics [Low] | -0.34 | (-)2.01-1.34 | 0.696 |
| Antibiotics [Median] | -0.35 | (-)2.03-1.33 | 0.683 |
| Observations | 1000 | | |
| R-squared/R-squared adjusted | 0.152 / 0.144 | | |

**Supplemental Table 8. Strain-specific effects of *B. vulgatus* monoculture growth in bile.**

| Predictors | Estimates | Confidence interval | p-value |
|---|---|---|---|
| (Intercept) | 8.42 | 7.97-8.86 | **<0.001** |
| Bile [Low] | -0.34 | (-)1.84-1.17 | 0.659 |
| Bile [Median] | -3.92 | (-)5.42-(-)2.41 | **<0.001** |
| Strain [b139H] | -0.1 | (-)0.72-0.51 | 0.737 |
| Strain [CFPLTA002-1B] | -0.09 | (-)0.71-0.52 | 0.764 |
| Strain [CFPLTA003-2B] | -0.04 | (-)0.65-0.58 | 0.908 |
| Strain [127M] | 0.17 | (-)0.44-0.79 | 0.579 |
| Strain [127O] | 0.07 | (-)0.54-0.69 | 0.812 |
| Bile [Low] x Strain [b139H] | 0 | (-)2.12-2.13 | 0.996 |
| Bile [Low] x Strain [CFPLTA002-1B] | 0.22 | (-)1.91-2.34 | 0.841 |
| Bile [Low] x Strain [CFPLTA003-2B] | -0.26 | (-)2.39-1.86 | 0.807 |
| Bile [Low] x Strain [127M] | -0.13 | (-)2.26-1.99 | 0.902 |
| Bile [Low] x Strain [127O] | -0.03 | (-)2.16-2.09 | 0.975 |
| Bile [Median] x Strain [b139H] | 4.58 | 2.46-6.71 | **<0.001** |
| Bile [Median] x Strain [CFPLTA002-1B] | 3.03 | 1.03-5.02 | **0.003** |
| Bile [Median] x Strain [CFPLTA003-2B] | 1.99 | (-)0.01-3.99 | 0.051 |
| Bile [Median] x Strain [127M] | 1.53 | (-)0.47-3.53 | 0.132 |
| Bile [Median] x Strain [127O] | 2.96 | 0.84-5.09 | **0.006** |
| Observations | 245 | | |
| R-squared/R-squared adjusted | 0.192 / 0.132 | | |

**Supplemental Table 9. Results from Mariner transposon (Tn*M*) screen in *E. coli* 139H.**

| | |
|---|---|
| **Colonies screened** | 9,858 (2x coverage) |
| **Non-viable *E. coli*** | 0 |
| **Primary hits (all)**<br>*Full restoration*<br>*Partial restoration*<br>*Minimal restoration* | 549 (5.56%)<br>87 (0.88%)<br>173 (1.75%)<br>289 (2.93%) |
| **Secondary hits**<br>*Full restoration*<br>*Partial restoration* | <br>51 (0.517% of total)<br>24 (0.243% of total) |
| **Mutants successfully sequenced**<br>*Full restoration*<br>*Partial restoration* | <br>47<br>20 |

**Supplemental Table 10. Mariner transposon mutants that allow for full and partial restoration of *B. vulgatus* viability in co-culture with glycerol.**

| "Full restoration" *E. coli* Tn*M* mutants (47/51 sequenced) | | | |
|---|---|---|---|
| **Category** | **Annotation** | **Function** | **Count** |
| Pathogenicity | *clb/pks* island (*clbBDJKN)* | Colibactin non-ribosomal peptide synthase | 26 (51%) |
| Metabolism | *actP* | Acetate symporter | 4 |
| | *dhaR* | Dihydroxyacetone kinase operon transcriptional regulator | 1 |
| | *malE* | Maltose binding protein | 1 |
| | *alsK* | Allose kinase | 1 |
| | *potF* | Spermidine/putrescine binding protein | 1 |
| | *gabP* | GABA permease | 1 |
| Stress | *sodA* | Superoxide dismutase | 1 |
| | *clpQY* promoter | Heat shock protein regulation | 1 |
| Biofilm/regulation | *pdeH* | Phosphodiesterase | 1 |
| | *rsmE* | rRNA methyltransferase | 2 |
| Phage | HKJJMI_25350 and HKJJMI_25825 | Tail sheath protein | 2 |
| | HKJJMI_25300 | Terminase, large sub-unit gp28 | 1 |
| rRNA | *5S/23S* | rRNA | 4 |
| **"Partial restoration" *E. coli* Tn*M* mutants (20/24 sequenced)** | | | |
| **Category** | **Annotation** | **Function** | **Count** |
| Pathogenicity | *htpG* | Molecular chaperone involved in colibactin synthesis | 2 |
| Biofilm/regulation | *rsmE* | rRNA methyltransferase | 5 |
| Metabolism | *pfkA* | 6-phosphofructokinase | 5 |
| | *menA* | 1,4-dihydroxy-2-naphthoate polyprenyltransferase | 3 |
| | *dhaK* | Dihydroxyacetone kinase subunit K | 1 |
| | *dhaM* | Dihydroxyacetone kinase phosphoryl donor | 1 |
| | *gldA* | Glycerol dehydrogenase | 1 |
| | *actP* | Acetate symporter | 1 |
| | *yijE* | Cysteine transporter | 1 |

**Supplemental Table 11. Mutants mapped to the *clb* island.** Attached separately due to size.

**Supplemental Table 12. Filtered blastp results of Clb proteins.** Attached separately due to size.
